# NuMA is a negative regulator of 53BP1 in DNA double-strand break repair

**DOI:** 10.1101/230706

**Authors:** Naike Salvador-Moreno, Jing Liu, Karen M. Haas, Laurie L. Parker, Chaitali Chakraborty, Stephen J. Kron, Kurt Hodges, Lance D. Miller, Paul J. Robinson, Sophie A. Leliévre, Pierre-Alexandre Vidi

## Abstract

Accumulation of 53BP1 at DNA breaks determines DNA repair pathway choice and promotes checkpoint activation. Here, we show regulation of 53BP1 beyond repair foci. 53BP1 movements are constrained in the nucleoplasm and increase in response to DNA damage. 53BP1 interacts with the structural protein NuMA, which controls 53BP1 diffusion. This interaction, and colocalization between the two proteins *in vitro* and in breast tissues, is reduced after DNA damage. In cell lines and breast carcinoma, NuMA prevents 53BP1 accumulation at DNA breaks and high NuMA expression predicts better patient outcomes. Manipulating NuMA expression alters PARP inhibitor sensitivity of BRCA1-null cells, end-joining activity, and immunoglobulin class switching that rely on 53BP1. We propose a new mechanism that involves the sequestration of 53BP1 by NuMA in the absence of DNA damage. Such mechanism may have evolved to disable repair functions and may be a decisive factor for tumor responses to genotoxic treatments.

## Introduction

DNA double-strand breaks (DSB) trigger a rapid and comprehensive DNA damage response (DDR) that leads to checkpoint signaling and cell cycle arrest, repair factor recruitment to the damage sites, and DNA repair. The precise orchestration of this response is critical for cell and organism survival (Ciccia & Elledge, 2010). Most DDR factors are permanent residents of the nucleoplasm that are not synthesized *de novo* during the DDR. Rather, repair foci formation relies on posttranslational modifications of histones and DDR factors (Price & D’Andrea, 2013). The DSB are processed predominantly by two competing pathways: Error-prone nonhomologous end-joining (NHEJ) of broken DNA strands, and homologous recombination (HR) that restores the genetic information from the sister chromatids. The committing step for HR is DNA end resection, whereas NHEJ is initiated by the recruitment of the KU70/KU80 scaffold at double-stranded DNA ends.

53BP1 is a key multifunctional DDR protein that plays an important role in repair pathway choice: 53BP1 and its effector RIF1 compete with BRCA1 to prevent CtIP-mediated resection (Callen *et al*., 2013, Chapman *et al*., 2013, Escribano-Diaz *et al*., 2013) and, as a consequence, antagonizes HR in favor of NHEJ. For DNA lesions undergoing HR repair, 53BP1 prevents excessive resection and thereby favors gene conversion over mutagenic single-strand annealing (Ochs *et al*., 2016). In the absence of functional BRCA1, the balance between HR and NHEJ is tilted, and DSB are improperly repaired by the NHEJ pathway, leading to deleterious chromosomal aberrations. This effect is exploited in anticancer therapies with PARP inhibitors (PARPi; reviewed in (Zimmermann & de Lange, 2014)). Acquired resistance limits clinical efficacy of PARPi, and loss of 53BP1 function is one of the mechanisms conferring PARPi tolerance in cancer cells (Bouwman *et al*., 2010, Bunting *et al*., 2010). With the exception of BRCA-null tumors, 53BP1 functions as a tumor suppressor, the loss of which radiosensitizes human (Squatrito *et al*., 2012) and mouse cells (Ward *et al*., 2003).

53BP1 is continuously expressed in the nucleus and rapidly accumulates at ionizing radiation-induced foci (IRIF) (Schultz *et al*., 2000). The recruitment of 53BP1 to IRIF requires binding of constitutive H4K20^Me2^ and of damage-induced H2AK15^ub^ marks by the tudor and ubiquitin-dependent recruitment (UDR) domains of the protein (Panier & Boulton, 2014). In addition, sustained 53BP1 function at IRIF depends on 53BP1’s BRCT domain binding to ATM-phosphorylated H2AX (Baldock *et al*., 2015). Much less is known about the regulation of 53BP1 spatial distribution and function outside of repair foci (if any). More generally, the mechanisms regulating repair factors access to chromatin in the absence of DNA damage remain largely unexplored. Yet such mechanisms may be key to prevent undue activation of the DDR. Here, we show that 53BP1 has a slow, non-Brownian nucleoplasmic diffusion behavior that accelerates in response to DNA damage. We identify a novel interaction between 53BP1 and the structural nuclear protein NuMA, which regulates the mobility, IRIF formation, and function of 53BP1 in DNA repair.

## Results

### 53BP1 movements are restrained in the nucleoplasm

The mechanisms that govern the spatial kinetics of 53BP1 in the absence of DNA damage are largely unknown. Therefore, we determined how the protein diffuses in regions of the nucleoplasm devoid of DNA breaks using fluorescence correlation spectroscopy (FCS) point measurements in cells expressing GFP-53BP1, and compared the values with predictions generated using an *in silico* diffusion model (Fig. 1A). Theoretical diffusion values for inert GFP molecules concurred with FCS measurements, which validated our model. The full structure of 53BP1 has not been solved; we therefore used hydrodiameters corresponding to sphere (8.2 nm) or rod (32.8 nm) structures to predict GFP-53BP1 diffusion. The FCS curves for GFP-53BP1 were best fitted with a two-component model interpreted as rapid Brownian motions coexisting with slower movements. The ‘Brownian’ diffusion values matched well with the predicted diffusion value assuming a spherical structure of 53BP1, which indicates that 53BP1 is likely to adopt a compact native conformation (Table S1). In contrast, the slower GFP-53BP1 motions differed largely from the predictions (Fig.1A). 53BP1 forms homo-oligomers *in situ*, independently of DNA damage (Adams *et al*., 2005). Yet, predicted diffusion values were significantly higher than FCS measurements (*P* < 0.0001), even when considering 53BP1 dimers or tetramers (Table S1). We therefore conclude that 53BP1 diffusion is constrained in the cell nucleus.

**Figure 1.**
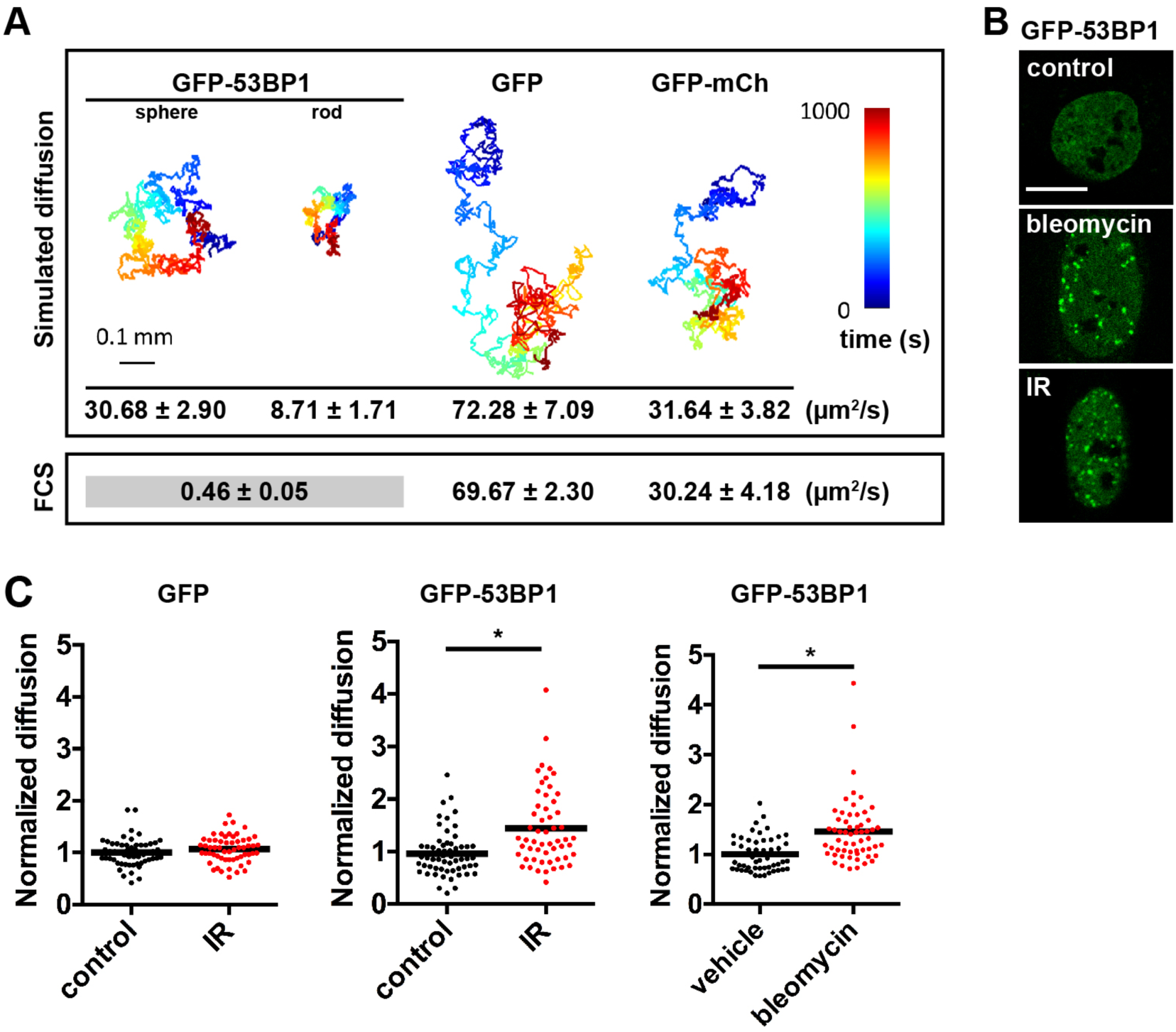
The diffusion of 53BP1 is constrained in the nucleoplasm and altered in response to DNA damage. **A** Modeling of GFP-53BP1 molecular dynamics in the viscoelastic context of the nucleus, assuming either a spherical or a rod-shaped structure of 53BP1. Simulated time traces and corresponding predicted diffusion times (mean ± SEM) are shown together with experimental values derived from FCS measurements (n ≥ 15 for simulation and FCS measurements). **B** Confocal images of U2OS cells expressing GFP-53BP1. Cells were left untreated (control), treated with bleomycin (20 mU/ml; 1h), or exposed to ionizing radiations (IR; followed by 30 min recovery). Scale bar, 10 μm. **C** Diffusion of GFP-53BP1 and GFP in cells treated as in **(B)**. The data represent measurements from at least 50 cells from two independent biological replicates. *, *P* < 0.0001 (t-test)

### DNA damage increases 53BP1 mobility

Restricted diffusion of 53BP1 may antagonize its recruitment to DNA breaks. To test whether the constraints on 53BP1 are modulated during the DDR, diffusion of GFP-53BP1 was measured in the absence and presence of DNA damage induced with ionizing radiations (IR) or with the radiomimetic drug bleomycin. As expected (Schultz *et al*., 2000), treatments with IR and bleomycin lead to the rapid focal accumulation of GFP-53BP1 (Fig. 1B). Although 53BP1 molecules are dynamically exchanged at repair foci (Pryde *et al*., 2005), the association of 53BP1 with the chromatin flanking DNA breaks strongly reduces 53BP1 mobility (Asaithamby & Chen, 2009), which was clearly apparent in FCS measurements (Fig. S1). To avoid this effect, we selected regions in-between repair foci for FCS point measurements. Exposures to IR and bleomycin both lead to a significant increase in GFP-53BP1 mobility outside repair foci (Fig. 1C). This increase could not be explained by altered nucleoplasm viscosity since the kinetics of GFP alone remained unchanged after DNA damage induction. Rather, the data suggest a mechanism regulating 53BP1 spatial kinetics in response to DNA damage.

### 53BP1 interacts with the nucleoskeletal protein NuMA

We used a proteomics approach to identify proteins interacting with 53BP1 and regulating its dynamics in the nucleus. Protein extracts derived from MCF-7 cells expressing a C-terminal portion of 53BP1 (53BP1_ct_; residues 1220-1711) fused to GFP were immunoprecipitated with GFP antibodies and analyzed by tandem mass spectrometry. This portion of 53BP1 was selected because the 53BP1 domains known so far to influence 53BP1 foci formation and oligomerization reside in the C-terminus of the protein, and full-length 53BP1 proved difficult to express. Among the proteins identified with the highest confidence (identification probability ≥ 95% and Mascot score > 200), 18.6% have a known function in genome maintenance and 9.9% have previously been identified as 53BP1 partners (Table S2). As expected, the proportion of known 53BP1-interacting proteins was much lower among the proteins identified with lower confidence. We shortlisted 53BP1 binding candidates identified in this study and in previous analyses of the 53BP1 interactome deposited in PubMed or in the BioGrid database (http://thebiogrid.org/) (Fig. 2A). Among these proteins, the nuclear mitotic apparatus protein (NuMA) was identified with the highest confidence in our analysis.

**Figure 2.**
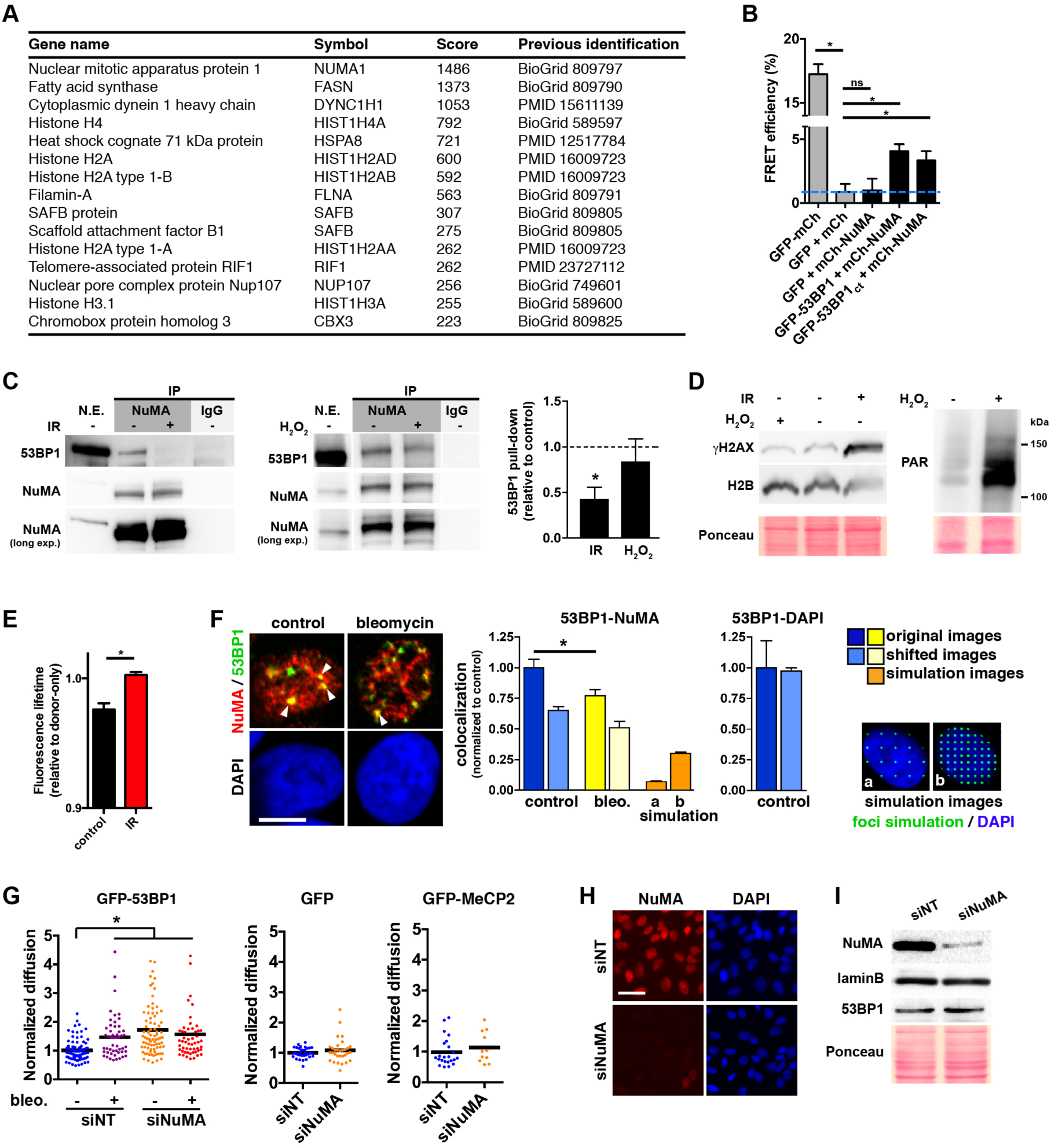
NuMA interacts with 53BP1 and regulates 53BP1 kinetics. **A** 53BP1-interacting partners identified by mass spectrometry in this and in previous studies. **B** Photobleaching FRET in U2OS cells expressing 53BP1 or 53BP1_ct_ and NuMA fused to GFP and mCherry (mCh). A GFP-mCh tandem construct was used as positive control, whereas coexpression of GFP and mCh was used as negative control. **C** Immunoprecipitation (IP) of NuMA from U2OS nuclear extracts (N.E.). Nonspecific immunoglobulins (IgGs) were used as controls. The western blots were probed for NuMA and 53BP1. Cells were exposed to IR (10 Gy, followed by 30 min recovery) or treated with H_2_O_2_ (1 mM, 10 min) prior IP. Densitometric quantification of 53BP1 pull down is shown in the graph. The ratio of 53BP1 over NuMA signals in IP samples was calculated and normalized to controls. The data represent mean ± SEM. *, *P* < 0.05 (one sample t-test; n = 4 (IR) or 3 (H_2_O_2_)). **D** Western blot analysis of γH2AX and PAR in the nuclear extracts used for IP. The results confirm the induction of DSB by IR and oxidative DNA damage by H_2_O_2_. Immunoblot for H2B and Ponceau staining are shown as loading controls. **E** FRET measured using fluorescence lifetime imaging in nonirradiated (control) and irradiated (IR) U2OS cells. 53BP1 and NuMA were labeled with antibodies coupled to Alexa Fluor 488 (FRET donor) or with Alexa Fluor 555 (FRET acceptor), respectively. Donor fluorescence lifetime was measured in the absence or presence of acceptor and used to calculate FRET efficiencies. Data represent mean ± SEM (n = 4). **F** Colocalization between 53BP1 and NuMA immunostaining in S1 cells treated with vehicle (control) or with bleomycin. Mander’s M2 colocalization coefficients for 53BP1 and NuMA or for 53BP1 and DAPI are shown in the bar graphs. Bars with paler colors quantify colocalization after shifting 53BP1 images by 10 pixels. Orange bars represent colocalization between NuMA and synthetic images with low (a) or high (b) foci density, as illustrated in the inset. Values were normalized to control. *, *P* < 0.05 (t-test, n = 12 image frames from three experiments). Scale bar, 5 μm. **G** FCS analysis of GFP-53BP1, GFP, and GFP-MeCP2 diffusion in cells transfected with nontargeting (NT) or with NuMA-specific siRNA. Cells were treated with bleomycin or vehicle. *, *P* < 0.001 (ANOVA and Tukey; n = 50–60 cells from two independent biological replicates). **H** NuMA silencing verified by immunostaining in U2OS cells transfected as in **(G)**. Scale bar, 50 μm **I** NuMA silencing verified by western blot with protein extracts from U2OS cells transfected as in **(G)**.

NuMA is a structural nuclear protein with a well-established function in spindle pole assembly and maintenance (Silk *et al*., 2009). In addition to mitotic functions, NuMA is essential for the establishment of higher-order chromatin organization during epithelial cell differentiation (Abad *et al*., 2007, Lelièvre *et al*., 1998) and for DNA repair by HR (Vidi *et al*., 2014). Fluorescence resonance energy transfer (FRET) experiments using the acceptor photobleaching method confirmed that mCherry-tagged NuMA interacts with GFP-53BP1 and GFP-53BP1_ct_. In contrast, low FRET efficiencies were measured when GFP alone was used as FRET donor (Fig. 2B). Co-immunoprecipitation (IP) experiments using NuMA antibodies confirmed the interaction between endogenous 53BP1 and NuMA in U2OS osteosarcoma cells, in HMT-3522 T4-2 breast cancer cells, and in non-neoplastic HMT-3522 S1 mammary epithelial cells (Fig. 2C; Fig. S2A-B). Importantly, the interaction between the two proteins decreased after DSB induction with ionizing radiations, but not after a short exposure to hydrogen peroxide used as a source of oxidative DNA damage (Fig. 2C-D). FRET measurements in cells labeled with fluorescent 53BP1 and NuMA antibodies confirmed the association between endogenous 53BP1 and NuMA in the absence of DNA damage and loss of interaction after IR (Fig. 2E).

To further address the link between NuMA and 53BP1, the relative distribution of the two proteins was quantified (Fig. 2F). NuMA signals overlapped with a subset of 53BP1 nuclear bodies in untreated cells. Shifting one of the two images by a few pixels decreased colocalization between 53BP1 and NuMA (but not between 53BP1 and DAPI), suggesting that the overlap was not random. Intriguingly, the colocalization between the two proteins decreased in cells exposed to bleomycin. By computing colocalization indices between NuMA and synthetic images with low or high foci densities, we determined that loss of colocalization between NuMA and 53BP1 in response to DNA damage was not due to an increase in the number of 53BP1 foci. 53BP1 nuclear bodies in nondamaged cells are distinct from 53BP1 IRIF induced by drugs and radiations and correspond to dysfunctional telomeres (Denchi & de Lange, 2007) as well as ‘shielded’ mitotic lesions in G1 cells (Lukas *et al*., 2011). The loss of colocalization and interaction of NuMA and 53BP1 after DNA damage induction therefore suggests that NuMA regulates 53BP1 function during the initial response to DNA damage.

### NuMA reduces 53BP1 mobility outside repair foci

The DNA damage-dependent interaction and colocalization between 53BP1 and the structural protein NuMA prompted us to examine whether NuMA regulates the dynamics of 53BP1 in the cell nucleus. To this end, the spatial kinetics of GFP-53BP1 at nuclear regions lacking damage foci was compared in cells transfected with nontargeting or with NuMA-targeting siRNA. GFP-53BP1 diffusion was significantly higher in cells silencing NuMA. This increase was similar to the increased diffusion resulting from DNA damage induction with bleomycin, and both treatments had no additive effect (Fig. 2G-I). Silencing NuMA did not alter the dynamics of GFP or GFP fused to the methyl CpG binding protein 2 (GFP-MeCP2), nor cause DNA damage (see next section). Hence, stabilization of 53BP1 diffusion measured in the absence of DNA damage requires NuMA.

### NuMA depletion does not alter 53BP1 expression

NuMA is structurally related to nuclear lamins, which have been shown to stabilize 53BP1 (Gibbs-Seymour *et al*., 2015, Gonzalez-Suarez *et al*., 2009, Mayca Pozo *et al*., 2017). In contrast to laminA/C depletion, silencing NuMA did not alter 53BP1 levels in U2OS or HMT-3522 S1 cells (Fig. 2I, Fig. 3A), consistent with previous observations (Vidi *et al*., 2014). Recent studies have shown that DNA damage can lead to 53BP1 degradation by the proteasome (Drane *et al*., 2017, Mayca Pozo *et al*., 2017). This effect was however not observed in U2OS or HMT-3522 S1 cells under our experimental conditions (Fig. S2C-D), which may be explained by different 53BP1 levels and distribution in soluble and insoluble nuclear compartments in the different cell lines (Mayca Pozo *et al*., 2017).

**Figure 3.**
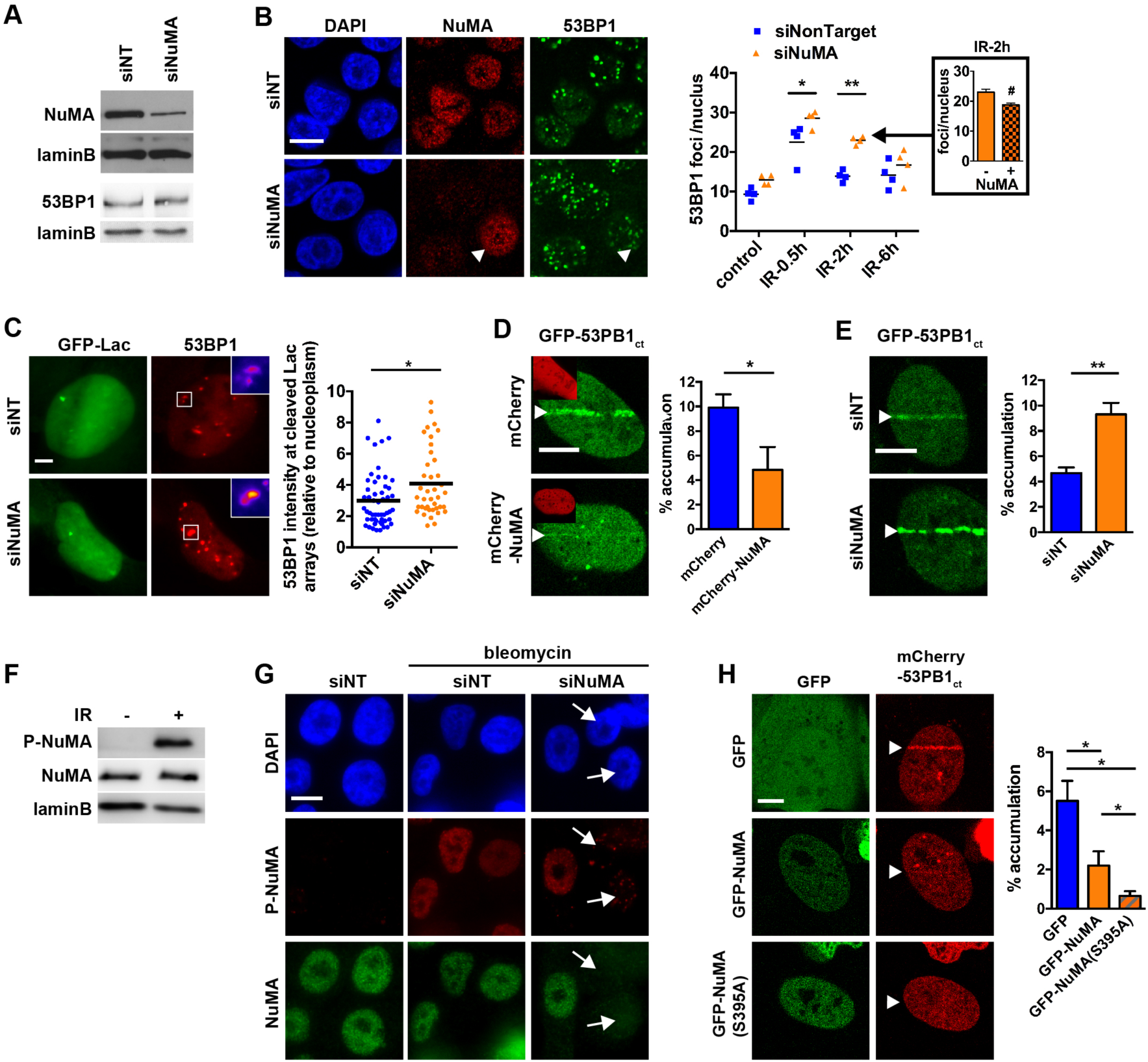
NuMA antagonizes 53BP1 accumulation at DNA damage sites. **A** NuMA silencing and 53BP1 expression verified by western blot. Lamin B is shown as loading control. **B** IR-induced focal accumulation of 53BP1 in HMT-3522 S1 cells transfected with nontargeting (NT) or NuMA siRNA. Confocal images show NuMA and 53BP1 immunostaining in irradiated cells (3 Gy) left to recover for 2h. The arrowhead points to a cell retaining NuMA expression in contrast to its neighbors. Quantification of 53BP1 foci density is presented in the bar graph. *, *P* < 0.05 and **, *P* < 0.001 (ANOVA and Tukey’s post-hoc test; n = 4). The inset shows the average number of 53BP1 foci per nucleus in cells silencing (−) or retaining (+) NuMA expression within the siNuMA transfection condition. #, *P* < 0.05 (t-test). **C** Relative intensity of 53BP1 immunostaining signals at cleaved ISceI sites. Cleavage sites were identified by labeling flanking Lac arrays with GFP-Lac. *, *P* < 0.01 (t-test; n ≥ 40 cells from three biological replicates). **D-E** Accumulation of GFP-53BP1_ct_ at laser-microirradiated tracks in U2OS cells expressing mCherry or mCherry-NuMA (insets; **D**) or transfected with nontargeting and NuMA siRNA **(E)**. The fractions of GFP signals at the tracks are shown in the bar graphs. * *P* < 0.05 and ** *P* < 0.0001 (t-test; n > 10 cells). **F** IR-induced NuMA phosphorylation at Ser395 (P-NuMA) in U2OS cells detected by immunoblot. **G** Localization of P-NuMA signals by immunostaining after bleomycin treatment (20 mU/ml; 1h) in cells transfected with NT or NuMA siRNA. **H** Accumulation of mCherry-53BP1_ct_ after laser-microirradiation in cells expressing GFP, GFP-NuMA, or a nonphosphorytable NuMA mutant [GFP-NuMA(S395A)]. * *P* < 0.05 (t-test; n > 15 cells from three biological replicates). Scale bars, 10 μm

### NuMA antagonizes 53BP1 recruitment at DNA breaks

To test if NuMA influences 53BP1 accumulation at DNA repair foci, HMT-3522 S1 mammary epithelial cells were transfected with NuMA-targeting or with nontargeting siRNA and exposed to IR. Increased 53BP1 foci density was measured in cells silencing NuMA compared to the nontargeting siRNA controls (Fig. 3A-B). Noticeably, within the NuMA siRNA transfections, irradiated cells retaining NuMA expression had significantly less 53BP1 foci compared to cells with no detectable NuMA expression. We could exclude that NuMA silencing sensitized the cells to IR since similar amounts of DSB were detected using the comet assay in siNuMA transfectants and controls (Fig. S3A). Moreover cell cycle distribution – an important factor in repair pathway choices – was not affected by NuMA depletion in S1 cells (Fig. S3B). Increased 53BP1 foci numbers was also measured in bleomycin-treated S1 cells expressing NuMA-targeting shRNA constructs (Fig. S3C) and a similar effect was measured for GFP-53BP1 in U2OS cells, where siRNA-mediated NuMA depletion lead to increased GFP-53BP1 foci formation in response to bleomycin or mitomycin C (Fig. S3D).

Next, we asked if NuMA affects 53BP1 accumulation at single DSB using a cell system with a stable integration of the ISceI cleavage site flanking lac arrays. In this system, DSB are induced and visualized by coexpression of ISceI and GFP fused to the lac repressor. 53BP1 foci detected by immunostaining at cleaved arrays were brighter in cells with NuMA silenced compared to controls (Fig. 3C). As an independent measure of 53BP1 recruitment at sites of DNA damage, U2OS cells expressing GFP-53BP1 were laser-microirradiated and the proportion of GFP fluorescence at the irradiation tracks was quantified. NuMA overexpression decreased GFP-53BP1 accumulation at the laser tracks and the same effect was observed for GFP-53BP1_ct_ (Fig. 3D and Fig. S4A). In contrast, silencing NuMA lead to increased accumulation of GFP-53BP1_ct_ at microirradiated lines (Fig. 3E). Overexpressing or silencing NuMA did not perturb PCNA recruitment to the laser tracks, ruling out nonspecific effects (Fig. S4B-C).

### Pan-nuclear NuMA phosphorylation regulates 53BP1 accumulation at sites of DNA damage

We showed previously that NuMA regulates SNF2h accumulation at DNA breaks (Vidi *et al*., 2014), yet silencing SNF2h did not alter 53BP1 line formation in microirradiation assays, suggesting that SNF2h does not mediate spatial regulation of 53BP1 by NuMA (Fig. S3D-E). These results are in agreement with a previous report showing no defect in 53BP1 IRIF formation in SNF2h-depleted cells (Smeenk *et al*., 2013).

NuMA is an ATM substrate that is rapidly phosphorylated at serine 395 in response to IR (Matsuoka *et al*., 2007, Vidi *et al*., 2014) (Fig. 3F). Immunostaining with P-NuMA (Ser 395) antibodies after bleomycin treatment or laser microirradiation revealed pan-nuclear NuMA phosphorylation in response to DNA damage (Fig. 3G and S5A), and specificity of the pan-nuclear P-NuMA signals was confirmed with human cells silencing NuMA and with mouse cells expressing human GFP-NuMA (Fig. 3G and S5B). These data indicate that NuMA is phosphorylated throughout the cell nucleus, i.e. not only at DNA damage sites. To determine if NuMA phosphorylation may regulate 53BP1 accumulation at damaged chromatin, we compared 53BP1 line formation after laser microirradiation in cells expressing GFP, GFP-NuMA, or GFP fused to a nonphosphorylatable NuMA mutant with Ser 395 substituted for an alanine [GFP-NuMA(S395A)]. Whereas GFP-NuMA expression reduced 53BP1 line accumulation by approximately 50%, expression of the nonphosphorylatable mutant almost completely abrogated 53BP1 recruitment to DNA damage sites (Fig. 3H). Western blot analysis confirmed phosphorylation of GFP-NuMA, but not GFP-NuMA(S395A), in response to radiations (Fig. S5C). Together, the data suggest that NuMA acts as a barrier preventing 53BP1 accumulation at damaged chromatin. NuMA phosphorylation by ATM may serve as a release mechanism.

### NuMA negatively regulates 53BP1 function in DNA repair

53BP1 is essential for immunoglobulin class switch recombination (CSR) during B cell maturation (Manis *et al*., 2004, Ward *et al*., 2004); a process where DSB induced by AID (activation-induced cytidine deaminase) in the immunoblobulin heavy chain gene are re-ligated by long-range NHEJ. First, we confirmed that silencing 53BP1 in CH12F3-2 murine B cells (Nakamura *et al*., 1996) reduces immunoglobulin switching to the IgA class (Fig. 4A). Next, we determined the effect of NuMA silencing and overexpression on CSR efficiency. NuMA depletion with shRNAs resulted in a modest (20%) but consistent increase in IgA switching, whereas GFP-NuMA overexpression led to a 60% reduction in CSR (Fig. 4A-B). Expression of the phospho-null NuMA mutant further exacerbated the CSR defect. Within samples nucleofected with the GFP empty vector, the proportion of IgA-switched cells was similar in GFP-positive and -negative cells (*P* > 0.05, paired t-test). In contrast, switching in GFP-positive vs. GFP-negative cells was reduced by 60% and 80% in the GFP-NuMA and GFP-NuMA(S395A) nucleofections, respectively (*P* = 0.0053 and *P* < 0.0001 relative to GFP).

**Figure 4.**
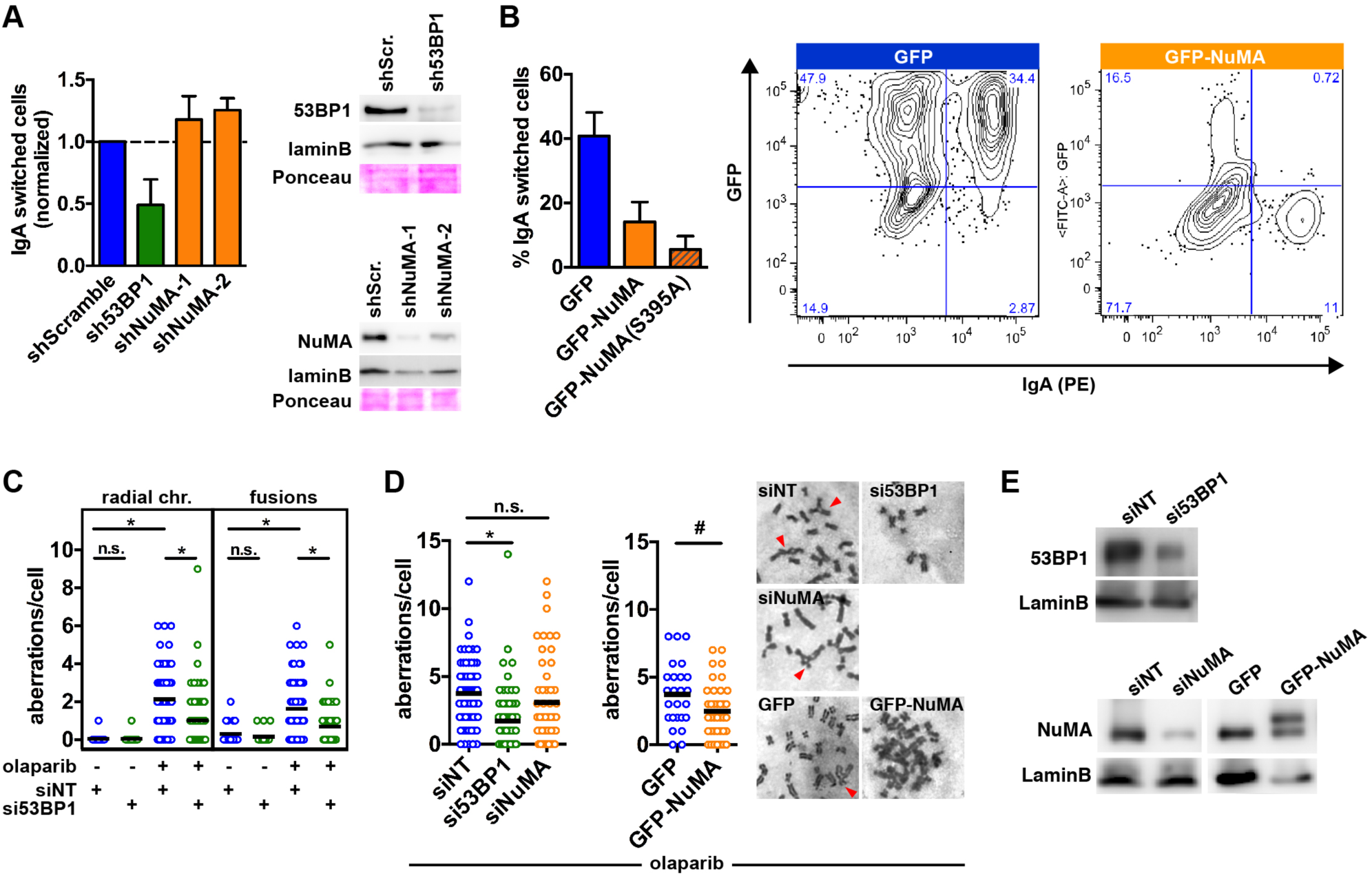
NuMA negatively regulates 53BP1 function in DNA repair. **A** Class switch recombination (CSR) in stimulated CH12F3-2 B cells stably transduced with scrambled, 53BP1, and NuMA-specific shRNA vectors. 53BP1 and NuMA silencing was verified by western blot. **B** CSR in cells nucleofected to express GFP, GFP-NuMA, or GFP-NuMA(S395A). CSR was quantified among GFP-positive cells. A representative flow cytometry analysis of cells expressing GPF and GFP-NuMA is shown on the right. **C** Quantification of radial chromosomes and chromosome fusions after vehicle or olaparib (0.5 μM; 24h) treatment of BRCA1-null SUM149 cells transfected with nontargeting or 53BP1 siRNA. *, *P* < 0.005 (ANOVA and Tukey, n ≥ 20). **D** Chromosomal aberrations (radials + fusions) scored in olaparib-treated SUM149 cells transfected with siRNAs, GFP, or GFP-NuMA. *, *P* < 0.005 (ANOVA and Tukey, n ≥ 50). #, *P* < 0.05 (t-test, n ≥ 25). Representative metaphase micrographs are shown. Arrowheads indicate aberrations. **E** Validation of 53BP1 and NuMA silencing and GFP-NuMA expression (upper band) in SUM149 cells by western blot.

In breast and ovarian tumors with BRCA1 mutations, loss of 53BP1 was shown to partially restore HR, thereby reducing PARP inhibitor toxicity (Bouwman *et al*., 2010, Bunting *et al*., 2010). As expected, siRNA-mediated depletion of 53BP1 in BRCA1-null SUM149PT breast cancer cells decreased the frequency of chromosomal aberration in response to the PARP inhibitor olaparib (Fig. 4C). While NuMA depletion did not alter olaparib sensitivity in these cells, GFP-NuMA overexpression significantly reduced olaparib efficacy, similarly to 53BP1 loss (Fig. 4D-E).

An important function of the [53BP1-RIF1] complex is to promote NHEJ by preventing DSB end-resection, the committing step for HR (Zimmermann & de Lange, 2014). Thus, we measured NHEJ activity after manipulating NuMA expression using a GFP reporter system (Mao *et al*., 2008) stably integrated in U2OS cells. In cells overexpressing mCherry-NuMA, NHEJ activity was decreased by 50% relative to controls expressing mCherry only (Fig. S6A). In addition, the proportion of NHEJ-competent (GFP-positive) cells with mCherry signals was significantly reduced in the mCherry-NuMA transfections compared to the mCherry transfection, although the transfection efficiencies for mCherry and for mCherry-NuMA were not different (32 ± 4% vs. 34 ± 4%, respectively). Conversely, NuMA silencing lead to higher NHEJ activity, independently of cell cycle alterations (Fig. S6B). Hence, NuMA counteracts the NHEJ pathway, consistent with its inhibitory effect on 53BP1 recruitment to DNA damage sites. From three independent assays (CSR, and PARPi sensitivity in the BRCA1-null context, and NHEJ efficacy), we conclude that NuMA negatively regulates 53BP1 function in DSB repair.

### NuMA expression predicts survival in breast cancer patients

To address the *in vivo* relevance of 53BP1 regulation by NuMA, invasive ductal carcinoma (IDC) tissue samples were collected after surgery, irradiated with a clinically-relevant dose of IR (3 Gy), and left to recover in medium for one or six hours. NuMA and 53BP1 labeled by immunofluorescence were detected by confocal microscopy (Fig. 5A). As expected, the number of 53BP1 nuclear foci increased in irradiated tissues compared to nonirradiated controls and subsequently decreased after the six-hours recovery period (Fig. 5B). Immunostaining confirmed the presence of NuMA in discrete nuclear spots in human epithelial tissues (Abad *et al*., 2007). Strikingly, the majority of the large 53BP1 nuclear bodies detected in nonirradiated cells overlapped with the NuMA spots. As observed in cell culture, colocalization between NuMA and 53BP1 decreased after IR (Fig. 5C). After recovery, the overlap between the two proteins went back to the levels found in the nonirradiated controls. We noticed and quantified large variations in NuMA expression within IDC samples (but not in normal breast tissues derived from reduction mammoplasties; Fig. S6A). This heterogeneity in NuMA levels enabled us to ask if NuMA expression affects 53BP1 foci formation. As observed *in vitro* (Fig. 3), NuMA levels negatively correlated with 53BP1 foci densities in irradiated IDC cells (Fig. 5D).

**Figure 5.**
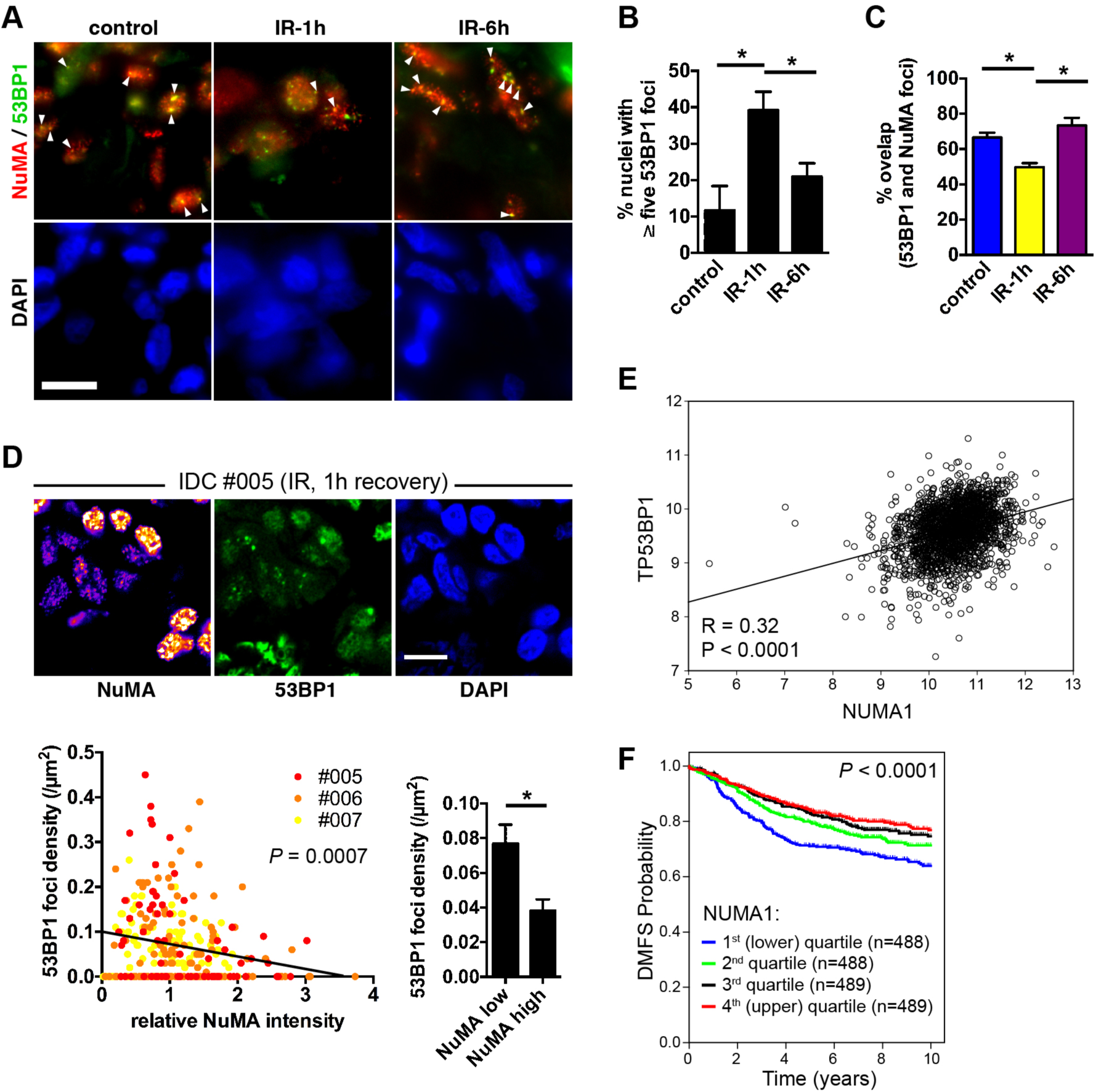
NuMA expression predicts survival in breast cancer patients. **A** Confocal images of 53BP1 and NuMA immunostaining in invasive ductal carcinoma (IDC) tissues. Tissue explants were mock-irradiated (control) or exposed to ionizing radiations (IR; 3 Gy) and left to recover for one or six hours. Nuclei were counterstained with DAPI. Arrowheads point to colocalized foci. **B** Quantification of 53BP1 foci in IDC tissues. **C** Overlap between 53BP1 foci and NuMA bright signals. *, *P* < 0.05 (ANOVA, tukey, n = 3). **D** Correlation between NuMA expression and 53BP1 foci in irradiated tissues (IDc patients #005-7). NuMA staining intensity is visualized with a heat map. For each cell nucleus, normalized NuMA intensities are plotted against the 53BP1 foci densities. The bar graph shows average foci densities in cells with low NuMA (first quintile) and high NuMA (last quintile). *, *P* < 0.005 (t-test). **E** Correlation analysis of *NUMA1* and *TP53BP1* mRNA expression levels in 1,954 breast tumors analyzed by microarray. Data are expressed as log2 normalized signal intensities. Pearson correlation (R) and p-value are shown. **F** Kaplan-Meier plot of distant metastasis-free survival (DMFS) with patients stratified by NUMA1 expression quartiles. The log-rank p-value is shown.

53BP1 is a potent tumor suppressor and loss of its expression drives tumorigenesis in multiple organs. On the other hand, 53BP1 repair function in cancer cells may promote cancer cell resistance to IR and chemotherapy (with the notable exception of BRCA1^-/-^ tumors) by enabling repair as well as by fueling genomic instability that results from nonhomologous end-joining. Indeed, increased radiosensitivity has been measured in 53BP1-deficient mice (Ward *et al*., 2003) and in glioblastoma tumors with very low levels of 53BP1 (Squatrito *et al*., 2012). A negative regulator of 53BP1 DDR functions may affect cancer cell survival in the presence of DNA damage and hence, cancer treatment responses. We therefore examined NuMA gene expression levels in a microarray dataset of breast tumor expression profiles. *NUMA1* transcript abundance was found to positively correlate with *TP53BP1* transcript levels (Fig. 5E) and to significantly associate with patient distant metastasis-free survival by Kaplan-Meier analysis (*P* < 0.0001). Stratification of tumors based on *NUMA1* messenger levels revealed a significant increase in survival for patients with tumors with high *NUMA1* expression (Fig. 5F). This effect persisted when only basal-like tumors were analyzed, indicating that the good prognosis associated with high *NUMA1* expression was not due to the enrichment of breast cancer subtypes with better outcomes among *NUMA1*-high cases (Fig. S6B). Association between *NUMA1* and *TP53BP1* expression was confirmed in an RNA-seq analysis of breast tumors (Fig. S6C). In this dataset, high *NUMA1* predicted longer overall survival (*P* < 0.0001; Fig. S6D).

## Discussion

We have shown that 53BP1 nucleoplasmic diffusion is at least one order of magnitude slower than theoretical values of free Brownian motions in the cell nucleus, irrespective of 53BP1 topology and multimerization parameters used for modeling. In cells with DNA damage, faster 53BP1 diffusion was measured in-between repair foci compared to nondamaged cells, which led us to hypothesize a nucleoplasmic retention mechanism in absence of DNA damage. The FCS method used for these analyses has two major advantages compared to bleaching approaches: Cells with low GFP-53BP1 signals could be analyzed, thereby avoiding overexpression artifacts, and nuclear regions could be precisely chosen for FCS point measurements, notably in-between IRIF. The results are consistent with biochemical fractionation experiments that detected 53BP1 in the insoluble nuclear fraction in the absence of DNA damage and in the chromatin fraction after IR (Gibbs-Seymour *et al*., 2015). A previous study determined the steady state kinetics of two other NHEJ factors, KU80 (14 μm^2^/s) and DNA-PKc (6 μm^2^/s) (Abdisalaam *et al*., 2014). These measurements are similar to the theoretical values for KU80 and DNA-PKc computed with our diffusion model (Table S1). In contrast to our finding with 53BP1, the diffusion of KU80 and DNA-PKc decreased after IR and bleomycin treatments (Abdisalaam *et al*., 2014), likely reflecting binding of the proteins to damaged chromatin regions. Reduced diffusion and chromatin immobilization after DNA damage was also demonstrated for SNF2h-containing ISWI chromatin remodeling complexes (Erdel *et al*., 2010). Therefore, the concept of repair factor sequestration that we put forward for 53BP1 may not apply for all DDR proteins. Rather, we postulate that this mechanism has evolved to ensure a tight control of key upstream DDR effectors such as 53BP1. Among its multiple DDR functions (Panier & Boulton, 2014), 53BP1 serves as a scaffold for the recruitment of DSB signaling and effector proteins, including EXPAND1/MUM1 that mediates chromatin decondensation in response to DNA damage (Huen *et al*., 2010). 53BP1 also amplifies ATM activity to promote checkpoint signaling (Panier & Boulton, 2014). While both aspects are essential for cell survival in the context of the DDR, it may be equally important for cellular homeostasis to prevent 53BP1 activation in the absence of DNA damage.

Our study identifies the structural nuclear protein NuMA as a binding partner and negative regulator of 53BP1. Interaction between the two proteins decreases in response to DNA damage and silencing NuMA enhances 53BP1 movement in the absence of DNA damage, suggesting that NuMA is a molecular component of the proposed retention mechanism. Accordingly, in cell lines and in breast cancer tissues, NuMA depletion increases 53BP1 accumulation at DNA damage sites while high NuMA levels leads to the opposite effect. ATM activity is both stimulated by 53BP1 and required for 53BP1 accumulation at IRIF, leading to a positive feedback loop after genomic insults and during physiological repair, including CSR; (Lumsden *et al*., 2004). Mechanistically, the UDR motif of 53BP1 binds to H2AK13/15 ubiquitinated by RNF168 downstream of ATM signaling (Fradet-Turcotte *et al*., 2013), and 53BP1’s BRCT domain binds to H2AX phosphorylated by ATM (Baldock *et al*., 2015), linking ATM activation to 53BP1 foci formation at damaged chromatin. Our results suggest an additional mechanism by which ATM controls 53BP1 retention by NuMA. Indeed, expression of NuMA(S395A) abolished 53BP1 recruitment at damage sites and blunted CSR activity.

Previously, we showed that NuMA targets the ISWI chromatin remodeler SNF2h to DNA breaks, thereby facilitating HR (Vidi *et al*., 2014). Several studies found that SNF2h activity is essential not only for HR, but also for NHEJ (Lan *et al*., 2010, Mueller *et al*., 2013, Smeenk *et al*., 2013). Surprisingly, NHEJ was not compromised in cells silencing NuMA. The experiments herein bring a molecular explanation: SNF2h loss-of-function in cells depleted from NuMA is likely counterbalanced by increased 53BP1 availability promoting NHEJ. It remains to be seen how generally NuMA and other structural elements of the cell nucleus regulate the DDR. Interestingly, NuMA associates with BRCA1, at least in the context of mitosis (Joukov *et al*., 2006). Moreover, other nuclear architectural factors have been implicated in the regulation of 53BP1: nucleoporin NUP153 promotes nuclear localization of 53BP1 (Lemaitre *et al*., 2012, Moudry *et al*., 2012), whereas A-type lamins regulate 53BP1 levels (Gibbs-Seymour *et al*., 2015, Gonzalez-Suarez *et al*., 2009), which accounts in part for genomic instability in laminopathies (Liu *et al*., 2005). In contrast, NuMA apparently does not facilitate nuclear import of 53BP1, nor regulates 53BP1 expression, but may rather sequestrate 53BP1 in the absence of damage to keep this master DDR effector in check. As such, NuMA functionally parallels the tudor interacting repair regulator (TIRR), a soluble factor preventing 53BP1 interaction with H4K20^Me2^ marks on the chromatin (Drane *et al*., 2017).

NuMA expression and localization is frequently altered in cancer cells (Bruning-Richardson *et al*., 2012, Knowles *et al*., 2006) and our analysis of protein and transcripts levels in breast tumors shows high levels of intra- and intertumor heterogeneity in NuMA levels, as well as a significant survival gain associated with high *NUMA1* gene expression in breast cancer patients. NuMA’s mitotic function may influence therapeutic responses. In addition, high NuMA levels in cancer cells may suppress the DSB repair function of 53BP1 – in particular mutagenic NHEJ linked to 53BP1 overexpression (Zong *et al*., 2015), thereby increasing tumor sensitivity to genotoxic anticancer treatments and lowering mutagenic rates that contribute to treatment resistances. These effects may be exploited for therapeutic purposes.

## Materials and Methods

### Cell culture, transfection, and genotoxic treatments

Osteosarcoma U2OS cells and HeLa cervical cancer cells were cultured in DMEM supplemented with 10% fetal bovine serum (FBS, Sigma). Non-neoplastic breast epithelial cells (HMT-3522 S1) were cultured in H14 medium (Vidi *et al*., 2013); HMT-3522 T4-2 breast cancer cells were cultured in H14 without EGF. SUM149PT breast cancer cells were cultured in DMEM supplemented with 10% FBS and with 10 mM HEPES buffer, hydrocortisone (5 μg/ml) and insulin (5 μg/ml). CH12F3-2 cells were cultured in RPMI 1640 containing 2 mM L-Glutamine, 10% FBS and 50 μM 2-mercaptoethanol in vertically positioned T25 flasks. Their concentration was kept below 10^5^ cells/ml. Lipofectamine 3000 (ThermoFisher) was used for siRNA (ON-TARGETplus, Dharmacon) and/or with plasmid DNA transfection. The following expression vectors were used for this study: pByffu (encoding a GFP-mCherry fusion protein) (Tramier *et al*., 2006); GFP-53BP1 and GFP-53BP1_ct_ (encoding full length 53BP1 and residues 1200 - 1711 of 53BP1 fused to GFP, respectively) (Huyen *et al*., 2004); mCherry-53BP1_ct_ (Addgene plasmid # 19835) (Dimitrova *et al*., 2008); GFP-Lac-NLS (Soutoglou *et al*., 2007); GFP-MeCP2 (Kudo *et al*., 2003); GFP-PCNA (Leonhardt *et al*., 2000); and mCherry-NuMA, cloned by replacing GFP in GFP-NuMA (Merdes *et al*., 2000) by mCherry using KpnI and BsrG1 restriction sites. GFP-NuMA(S395A) was cloned by overlap PCR using the 5’CAGCTGGAAGAACACcTtgCgCAGCTGCAGGATAACCCAC 3’ and 5’GTGGGTTATCCTGCAGCTGCGcAAGGTGTTCTTCCAGCTG 3’ primer pair. The overlap PCR product was digested with EcoRV and AflII and the fragment (1166 bp) was ligated into the corresponding sites of pcDNA GFP-NuMA. Clones were verified using restriction for presence of the FspI site (introduced by silent mutagenesis) and by DNA sequencing. The shRNA vectors targeting human NuMA were purchased from Origene (TR311065). The shRNA vectors for murine proteins were purchased from the Dharmacon RNAi Consortium and included shNuMA-1 (TRCN0000072130), shNuMA-2 (TRCN0000072132), and sh53BP1 (TRCN0000081778). A shRNA scramble pLKO.1 plasmid (Addgene # 1864) (Sarbassov *et al*., 2005) was used as negative control. Lentiviral particles generated using HIV packaging were used for transduction. Stably silenced cell lines were generated after selection with puromycin (0.6 μg/ml). DNA damage was induced by gamma irradiation (3 Gy and 10 Gy for S1 and U2OS cells, respectively; Gammacell 220 irradiator from Nordion), with the radiomimetic drug bleomycin (20 mU/ml for 1h), with MMC (2.6 μM, 18h), or with hydrogen peroxide (1 mM, 10 min).

### Modeling and measurements of protein diffusion

Simulations of Brownian motions were performed in Matlab and the diffusion coefficients were deduced from the mean squared displacement values of the simulations. All simulations were performed at 37°C. More details on the modeling approach can be found in the Expanded Experimental Procedures. Experimental diffusion coefficients were derived from FCS measurements. All the FCS experiments were performed on a customized scanning confocal microscope (Microtime 200, PicoQuant; (Liu & Irudayaraj, 2013)), as described previously (Vidi *et al*., 2014). Cells were maintained 37°C and 5% CO_2_ using a stage-top incubator (Tokai Hit).

### Mass spectrometry

GFP-fused to 53BP1 containing amino acids 1-34 and 1220-1711 (Huyen *et al*., 2004) was stably expressed in MCF7 cells under the Tet-On promoter. Cells were exposed to doxycycline (1 μg/ml) for 72 h to induce expression of the GFP-53BP1 construct. Cell lysates were immunoprecipitated using GFP antibodies, resolved by SDS-PAGE, and subjected to trypsin digestion. Peptides were separated by HPLC and analyzed by LC-MS/MS using a LTQ Orbitrap XL (Thermo). Spectra were extracted by ProteoWizard and were analyzed using Mascot (Matrix Science). Scaffold (Proteome Software Inc.) was used to validate MS/MS based peptide and protein identifications. More information on the mass spectrometry analysis can be found in the Expanded Experimental Procedures.

### Immunoprecipitation and western blot analysis

For coimmunoprecipitation, nuclear extracts were prepared using the Universal Magnetic Co-IP kit (Active Motif) in the presence of protease inhibitors (aprotinin, 10 μg/ml, Sigma; Pefabloc, 1 mM, Roche Applied Science), phosphatase inhibitor (sodium fluoride, 250 μM), and PARG inhibitor (DEA,10 μM, Trevigen). Nuclear extracts (0.5 mg) were immunoprecipitated with NuMA antibodies or nonspecific IgG (1.5 μg) overnight at 4°C, and analyzed by immunoblotting. Similar results were obtained with two different NuMA antibodies (Calbiochem, ab-2 and Bethyl Laboratories, A301-509A). Antibodies used for immunoblot were 53BP1 (Abcam, Ab36823, 1 μg/ml), γH2AX (Ser139; Millipore, clone JBW301, 1 μg/ml), Histone H2B (Abcam, Ab1790, 0.1 μg/ml), lamin B (Abcam, Ab16048, 60 ng/ml), NuMA (B1C11, 1:2, a gift from Dr. Jeffrey Nickerson, UMass, Worcester, USA), P-NuMA (S395; Cell Signaling, 3429, 1:1000), and PAR (Trevigen, 4336-APC-050, 1:1000).

### Immunostaining, laser microirradiation and FRET

Immunostaining was performed as described (Vidi *et al*., 2014). Antibodies used for immunofluorescence were 53BP1 (Abcam, 5 μg/ml), γH2AX (Millipore, 2 μg/ml), NuMA (B1C11, 1:2 or Abcam, clone EP3976, 1:250), P-NuMA (Cell Signaling, 1:200), and SNF2h (Abcam Ab3749, 20 μg/ml). Fluorescent signals were imaged with a Zeiss CLSM710 confocal microscope using a 63x oil (NA = 1.4) objective. Repair foci were quantified using a custom macro in ImageJ (http://rsbweb.nih.gov/ij/). For FLIM-FRET experiments, cells were fixed and stained with 53BP1 and NuMA antibodies labeled with Alexa Fluor^®^ 488 and −555 dyes, respectively. Fluorescence lifetime was collected and analyzed as previously (Vidi *et al*., 2014). For live cell imaging, cells in glass-bottom dishes (MatTek) were maintained at 37°C and 5% CO_2_ with a stage-top incubator (Pecon). The 405 nm laser line of the CLSM710 was used at maximum power to microirradiate lines across nuclei (100 μs dwelling time; ~1 s scan/line). Cells were sensitized with BrdU (10 μM) 48h prior microirradiation and were irradiated and imaged with a 63x water immersion objective (NA = 1.2) 5 min after microirradiation. For acceptor photobleaching FRET experiments, mCherry was bleached in spot areas of the cell nucleus using three pulse iterations (561 diode laser at 100% power and 177 μs/pixel dwell time). No FRET was detected in nonbleached regions.

### GFP-based repair assays and cell cycle analysis

NHEJ was assessed in U2OS cells with a stable integration of the NHEJ-I reporter cassette, as described (Mao *et al*., 2008). GFP-positive cells were quantified by flow cytometry. 10,000 – 100,000 cells were analyzed in each sample. Cells were costained with DRAQ5 (Biostatus) to collect DNA content values for cell cycle determination.

### Class switch recombination (CSR)

CH12F3-2 cells were nucleofected with shRNA vectors and overexpression plasmids (Lonza Nucleofector™ I device; solution L; program T20; 10^6^ cells and 2 μg DNA per reaction). CSR to IgA was induced by stimulating the cells for 72h with 5 μg/ml agonist anti-CD40 (clone HM40-3; BD), 5 ng/ml IL-4 (R&D Systems) and 2.5 ng/ml TGF-β1 (R&D Systems). Surface expression of IgA was analyzed by flow cytometry using PE-conjugated goat antimouse IgA antibodies (1:400; SouthernBiotech).

### Quantification of chromosomal aberrations

SUM149PT cells were seeded in 6-well plates (one plate per condition) and cultured until they reached 60% confluency for transfection. GFP and GFP-NuMA overexpressing cells were selected with geneticin (200 μg/ml). To quantify chromosomal aberrations, the cells were transferred onto 22×22 mm coverslips in 35 mm dishes 48h after transfection. Cells were then treated with PARP inhibitor (olaparib; 0.5 μM) or vehicle. After 24h incubation, colcemid (10 μg/ml) was added to the cells for 1h. Metaphase spreads were prepared by hypotonic bursting in (2:1) (75 mM KCl: 0.8% sodium citrate) and fixation in (3:1) (methanol: acetic acid). Subsequently, coverslips were air dried, mounted on microscope slides, stained with 2% Giemsa for 3 min and washed in Gurr buffer. Slides were allowed to dry and were analyzed with an Olympus IX83 microscope using a 60 X oil objective and CMOS camera (0RCA-Flash4.0; Hamamatsu).

### Human breast tissues

Breast invasive ductal carcinoma tissues were obtained from mastectomies and normal breast tissue was obtained from reduction mammoplasties performed at the IU Health Arnett Hospital (Lafayette, IN), after obtaining patient consent (Purdue Institutional Review Board approval 1206012467). The excision specimen were resected, minced to approx. 0.5 cm fragments, and placed in RPMI within 30 min of surgery. Tissue explants were rinsed with, and transported in RPMI. After (mock) irradiation (3 Gy; Gammacell 220 irradiator) and recovery in a tissue culture incubator (37°C; 5% CO_2_), tissue explants were incubated in PBS containing 18% sucrose on ice for 15 min, then with PBS with 30% sucrose on ice for 15 min, and frozen in optimal cutting temperature (OCT) compound on dry ice mixed with 70% ethanol. Tissues were stored at −80°C until sectioning of 12 μm thick sections with a cryostat set at −20°C. Tissue slices were collected on Superfrost plus slides and used for immunostaining.

### Analysis of Tumor Expression Profiles

Breast tumor expression profiles generated by The Cancer Genome Atlas (TCGA) Research Network (http://cancergenome.nih.gov/) and the microarray dataset of Nagalla et al. (Nagalla *et al*., 2013) were utilized in this study. The TCGA clinical data and RNA-seq data (level 3-processed) were downloaded from the Broad’s FireBrowse website (http://firebrowse.org/; TCGA data version 2016_01_28). The RNA-seq breast tumor data comprising 1,100 tumor expression profiles was filtered to exclude male and gender ‘unknown’ samples (n=13), metastatic tissue samples (n=7) and one errant skin cancer sample yielding 1,079 female primary breast tumor samples. OncoLnc (http://www.oncolnc.org) was used to visualize overall survival (Anaya, 2016). The dataset of Nagalla et al. was utilized as previously described (Nagalla *et al*., 2013).

## Acknowledgements

We are grateful to C. Cardoso, M. Coppey-Moisan, V. Gorbunova, T. Halazonetis, T. de Lange, T. Misteli, D. Sabatini, and J. Nickerson for generously sharing antibodies and DNA constructs, and to E. Alli and T. Honjo for providing the SUM149PT and CH12F3-2 cell lines. We thank M. Pettenati for assistance in the analysis of chromosomal aberration, K. Ragheb, J. Sturgis, M. Gray and J. Maldonado for technical help, and J. Muhlemann for discussions and helpful comments on earlier versions of the manuscript. This work was supported by the National Institutes of Health (K99/R00CA163957 to P.-A.V. and R01CA112017 to S.A.L.), by the American Cancer Society (RSG-12-170-01-LIB to K.M.H.), by the Wake Forest Center for Molecular Signaling (CMS) via its imaging facility, and by the National Cancer Institute’s Cancer Center Support Grant award number P30CA012197 issued to the Wake Forest Baptist Comprehensive Cancer Center. The content is solely the responsibility of the authors and does not necessarily represent the official views of the National Cancer Institute

## Author Contributions

NSM: photobleaching FRET, laser microirradiation, CSR assays, and quantification of chromosomal aberrations. JL: FCS, FLIM-FRET, and diffusion modeling. KMH: CSR assays. LLP, CC, SJK: mass spectrometry. KH: human breast tissue procuration. LDM: breast tumor expression profiles. PJR: assistance with flow cytometry. PAV: immunoprecipitations, colocalization analysis, NHEJ assays, and all other experiments. SAL, PAV: supervised the study. PAV, wrote the manuscript.

## EXPANDED EXPERIMENTAL PROCEDURES

### Modeling 53BP1 diffusion in the nucleus

Simulations of protein diffusion in the nucleus were implemented in Matlab, Macromolecules in solution undergo random collisions from surrounding molecules, resulting in a three dimensional (3D) translational or rotational “random walk”. According to Einstein’s theory, 3D translational dynamics in medium satisfies the diffusion equation:

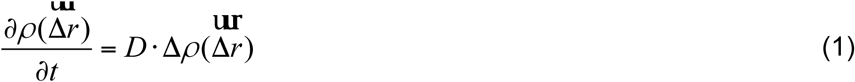

where 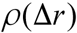 is the displacement distribution of the molecule, *D* is the diffusion coefficient, and Δ is the 3D Laplace operator. Theoretically, the diffusion coefficient has the relationship with the size, *d*, of the macromolecules

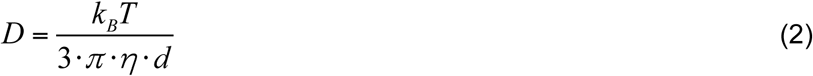

with *k_B_*, the Boltzmann constant, *T* the temperature, *η* the viscosity of the medium (1.5× 10^−3^ Pa-s in the nucleus (Liang, Wang et al., 2009).

The diffusion coefficient can also be obtained from mean squared displacement (MSD) calculations, 〈(Δ*r*)^2^〉. In our model, we only considered the Brownian motion of the protein in a 2D plane. Therefore,

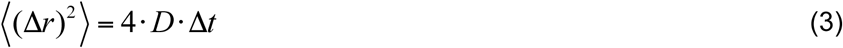

With Δ*t*, the lag time between each position Δ*r*.

We simulated the 2D Brownian motions of the proteins in time windows of 1000 s and the trajectories (*x_i_*, *y_i_*) per second (Δ_*t*_) were registered for calculation. The displacements were then calculated as:

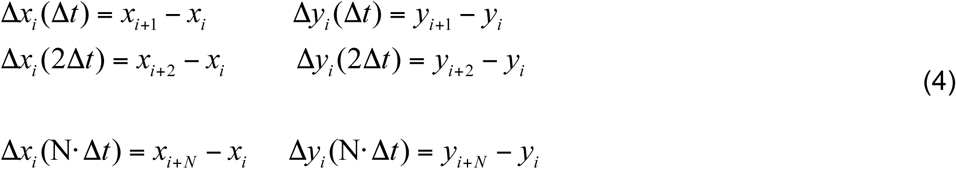

And the square displacement (Δ*r*)2 summarized as:

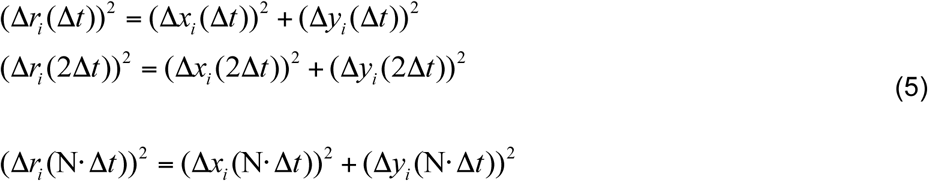

The diffusion coefficients *D* depend on the hydrodiameter (hence the shape) of the molecules (equation 2). GFP has a beta barrel structure of 4.2 nm in length (Ormo, Cubitt et al., 1996), used for modeling. To simulate GFP-53BP1 diffusion, we considered two extreme possibilities, a single globular domain (sphere) and a highly elongated molecule (rod). The corresponding physical sizes were predicted using a sedimentation model described in (Erickson, 2009). A sphere the size of GPF-53BP1 would have a diameter of 8.22 nm, whereas the calculated length of the rod would be 32.88 nm (using a sedimentation coefficient *S_max_/S* = 4). When predicting the sizes of 53BP1 dimers and tetramers, we considered combinations of spheres with minimum surface (16.44 nm and 19.84 nm for the dimer and tetramer, respectively), as well as concatenated rods (65.56 nm and 131.12 nm for dimers and tetramers, respectively). The diffusion coefficients were obtained by fitting MSD curves using equation (3).

### Mass spectrometry

Cells were then treated with paraformaldehyde (0.05% w/v) to cross-link proteins, lysed with lysis buffer (1% SDS, 10mM EDTA, 50 mM Tris-HCl pH 8.0) supplemented with Protease Inhibitor Cocktail (Roche) on ice, using probe sonication (also on ice) to shear the chromatin. GFP-53BP1 and associated proteins were immunoprecipitated from the lysate using μMACS anti-GFP conjugated magnetic microbeads (Miltenyi Biotech). Proteins were eluted and cross-links reversed by heating the beads in Laemmli loading buffer at 95°C for 5 min, followed by separation via SDS-PAGE on a 4-12% gradient gel (Invitrogen).

#### Trypsin Digestion of Samples from SDS-PAGE Gels

Lanes from 4-12% MOPS buffer SDS-PAGE gels were excised and cut into four equally-sized pieces using a sterile razor blade and chopped into ~1 mm^3^ pieces. Each sample was washed in water and destained using 100 mM ammonium bicarbonate pH 7.5 in 50% acetonitrile. A reduction step was performed by addition of 100 μl 50 mM ammonium bicarbonate pH 7.5 and 10 μl of 10 mM Tris(2-carboxyethyl)phosphine HCl at 37°C for 30 min. The proteins were alkylated by adding 100 μl 50 mM iodoacetamide and allowed to react in the dark at 20°C for 30 min. Gel samples were washed in water, then acetonitrile, and dried in a SpeedVac. Trypsin digestion was carried out overnight at 37°C with 1:50 enzyme-protein ratio of sequencing grade-modified trypsin (Promega) in 50 mM ammonium bicarbonate pH 7.5, and 20 mM CaCl_2_. Peptides were extracted with 5% formic acid and vacuum dried.

#### HPLC for Mass Spectrometry

The peptide samples from trypsin digestion were loaded to a 0.25 μl C_8_ OptiPak trapping cartridge custom-packed with Michrom Magic C8 (Optimize Technologies), washed, then switched inline with a 20 cm by 75 μm C_18_ packed spray tip nano column packed with Michrom Magic C18AQ, for a 2-step gradient. Mobile phase A was water/acetonitrile/formic acid (98/2/0.2) and mobile phase B was acetonitrile/isopropanol/water/formic acid (80/10/10/0.2). Using a flow rate of 350 nl/min, a 90 min, 2-step LC gradient was run from 5% B to 50% B in 60 min, followed by 50%–95% B over the next 10 min, hold 10 min at 95% B, back to starting conditions and re-equilibrated.

#### LC-MS/MS Analysis

The samples were analyzed via electrospray tandem mass spectrometry (LC-MS/MS) on a Thermo LTQ Orbitrap XL, using a 60,000 RP survey scan, m/z 375-1950, with lockmasses, followed by 10 LTQ CID scans on doubly and triply charged-only precursors between 375 Da and 1500 Da. Ions selected for MS/MS were placed on an exclusion list for 60 seconds.

#### Database searching

Tandem mass spectra were extracted by ProteoWizard version 3.0.3402. Charge state deconvolution and deisotoping were not performed. All MS/MS samples were analyzed using Mascot (Matrix Science, London, UK; version 2.4.1). Mascot was set up to search the uniprot_human_050713 database (selected for Homo sapiens, version downloaded May 7, 2013, 70555 entries) assuming the digestion enzyme trypsin. Mascot was searched with a fragment ion mass tolerance of 1.00 Da and a parent ion tolerance of 10.0 PPM. Oxidation of methionine, formyl of the n-terminus, carbamidomethyl of cysteine and phospho of serine, threonine and tyrosine were specified in Mascot as variable modifications.

#### Criteria for protein identification

Scaffold (version Scaffold_4.4.5, Proteome Software Inc., Portland, OR) was used to validate MS/MS based peptide and protein identifications. Protein probabilities were assigned by the Protein Prophet algorithm (Nesvizhskii, Keller et al., 2003). Peptide identifications were accepted if they could be established at greater than 95.0% probability by the Peptide Prophet algorithm (Keller, Nesvizhskii et al., 2002). Proteins that contained similar peptides and could not be differentiated based on MS/MS analysis alone were grouped to satisfy the principles of parsimony. Proteins were annotated with GO terms from NCBI (downloaded Jul 30, 2014) (Ashburner, Ball et al., 2000).

### EXPANDED VIEW TABLE AND FIGURES

**Table S1.**
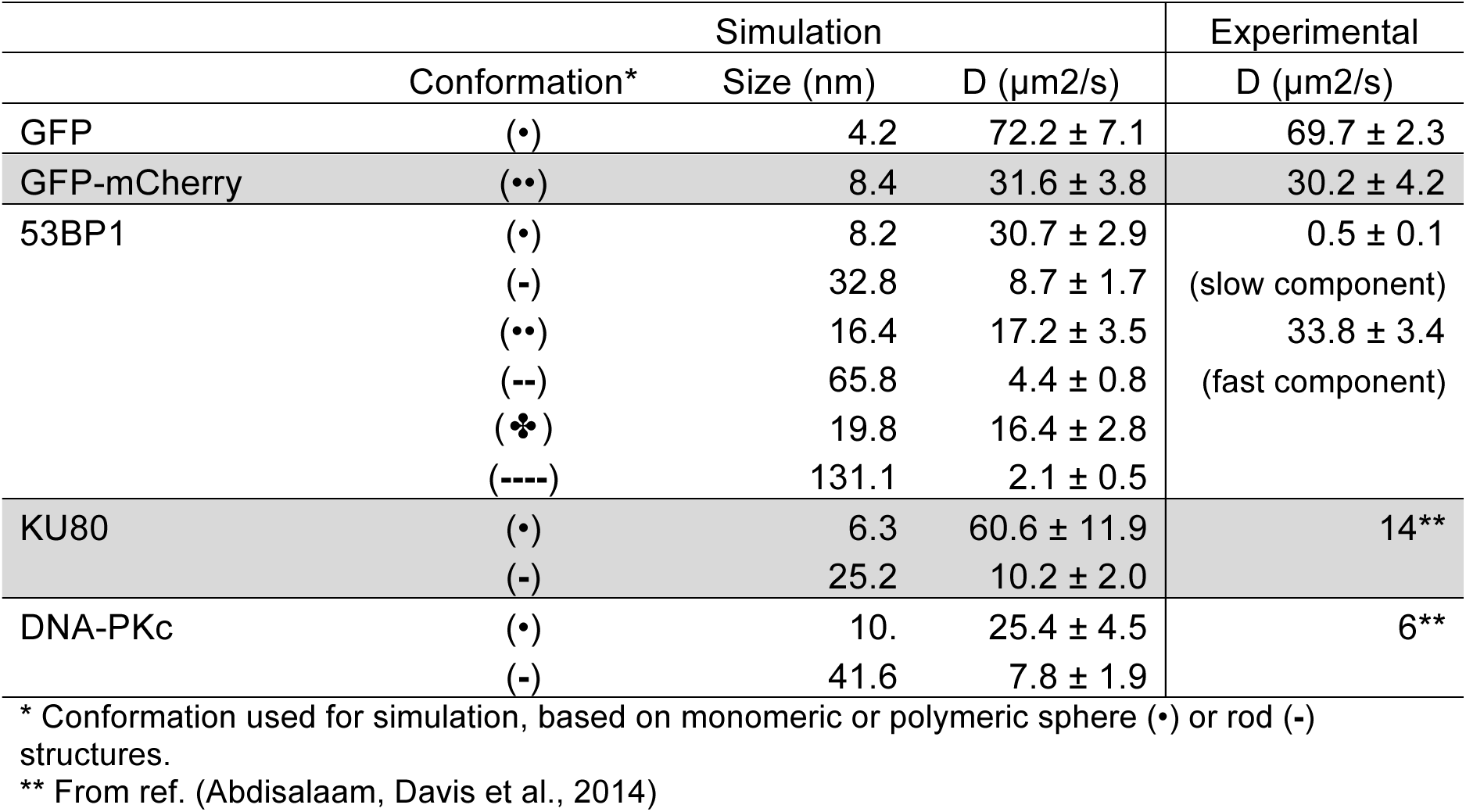
Predicted and measured diffusion times in the cell nucleus

**Table S2.**
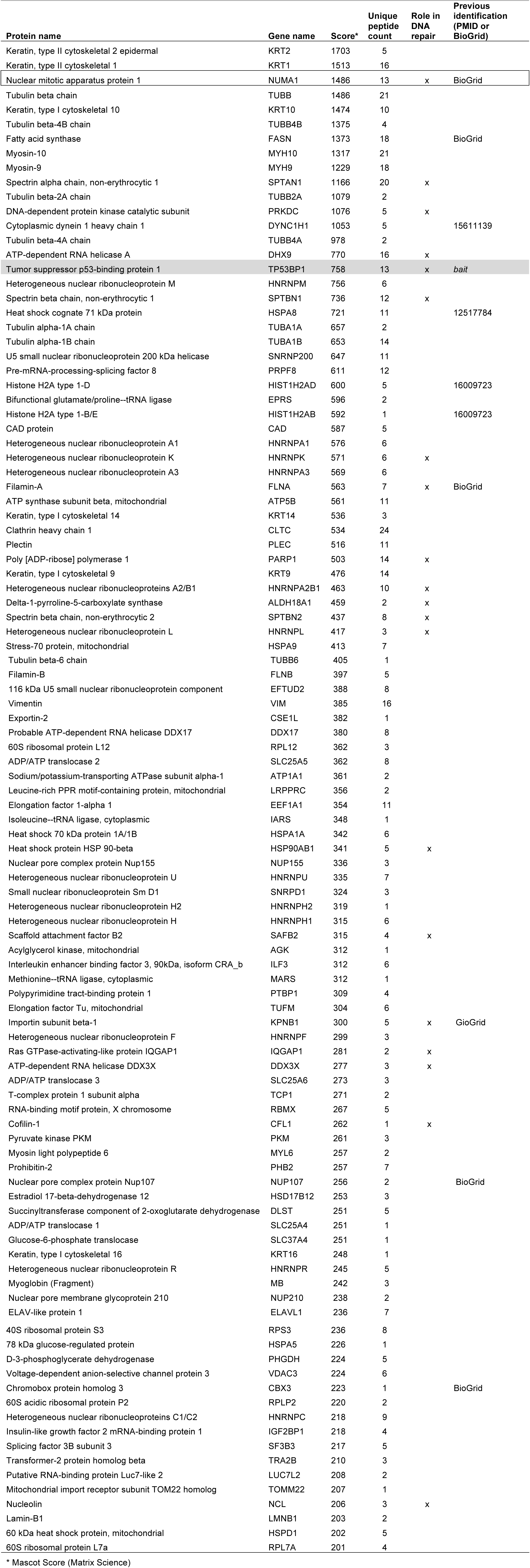
53BP1-binding proteins identified by mass spectrometry.

**Figure S1.**
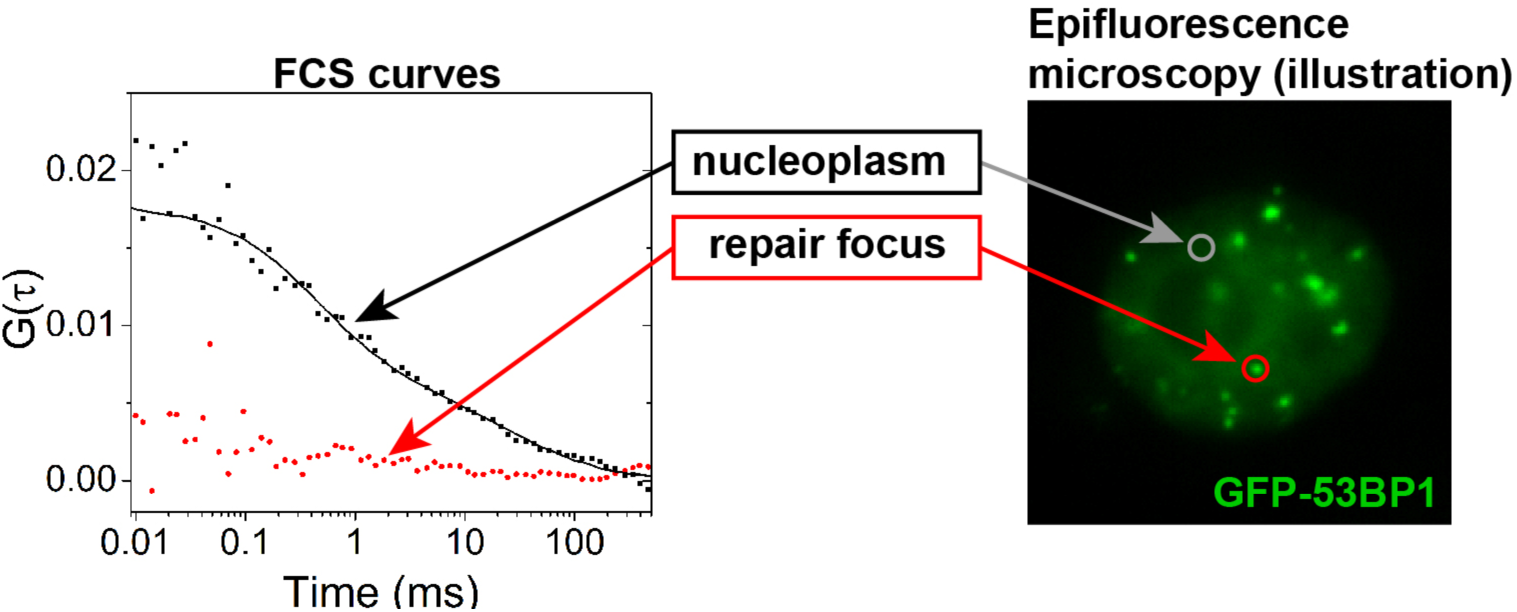
Representative FCS traces of GFP-53BP1 measured in the nucleoplasm in-between repair foci or at a repair focus in a bleomycin-treated cells. The flat autocorrelation curve corresponding to the measurement in a repair focus reflects slow diffusion kinetics.

**Figure S2.**
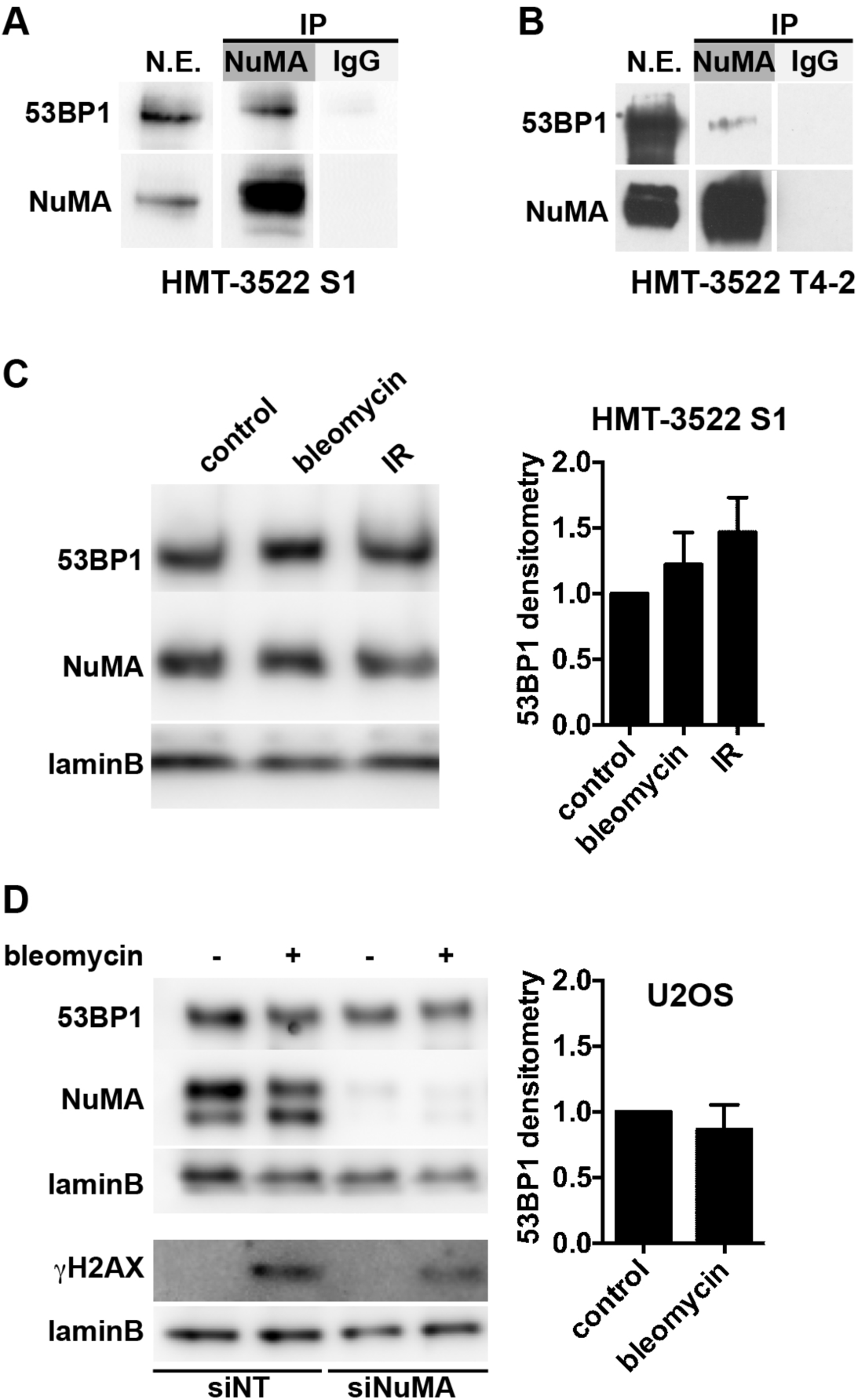
53BP1 interaction with NuMA and 53BP1 abundance after DNA damage. **A-B** Immunoprecipitation of NuMA from non-neoplastic HMT-3522 S1 **(A)** and malignant HMT-3522 T4-2 **(B)** nuclear extracts. Nonspecific IgGs were used as control. The blots were probed for NuMA and 53BP1. **C** 53BP1 levels in HMT-3522 S1 cells untreated (control), treated with bleomycin (20 mU/ml, 1h), or exposed to ionizing radiations (IR, 3 Gy followed by 1h recovery). Densitometry analysis of 53BP1 levels is shown in the bar graph (mean ± SEM; n = 4). **D** 53BP1 levels in U2OS cells after bleomycin treatment. Cells were transfected with NuMA (siNuMA) or nontargeting (siNT) siRNA. The γH2AX and NuMA immunoblots verify DNA damage induction and silencing, respectively. Densitometry analysis of 53BP1 levels is shown in the bar graph (mean ± SEM; n = 4).

**Figure S3.**
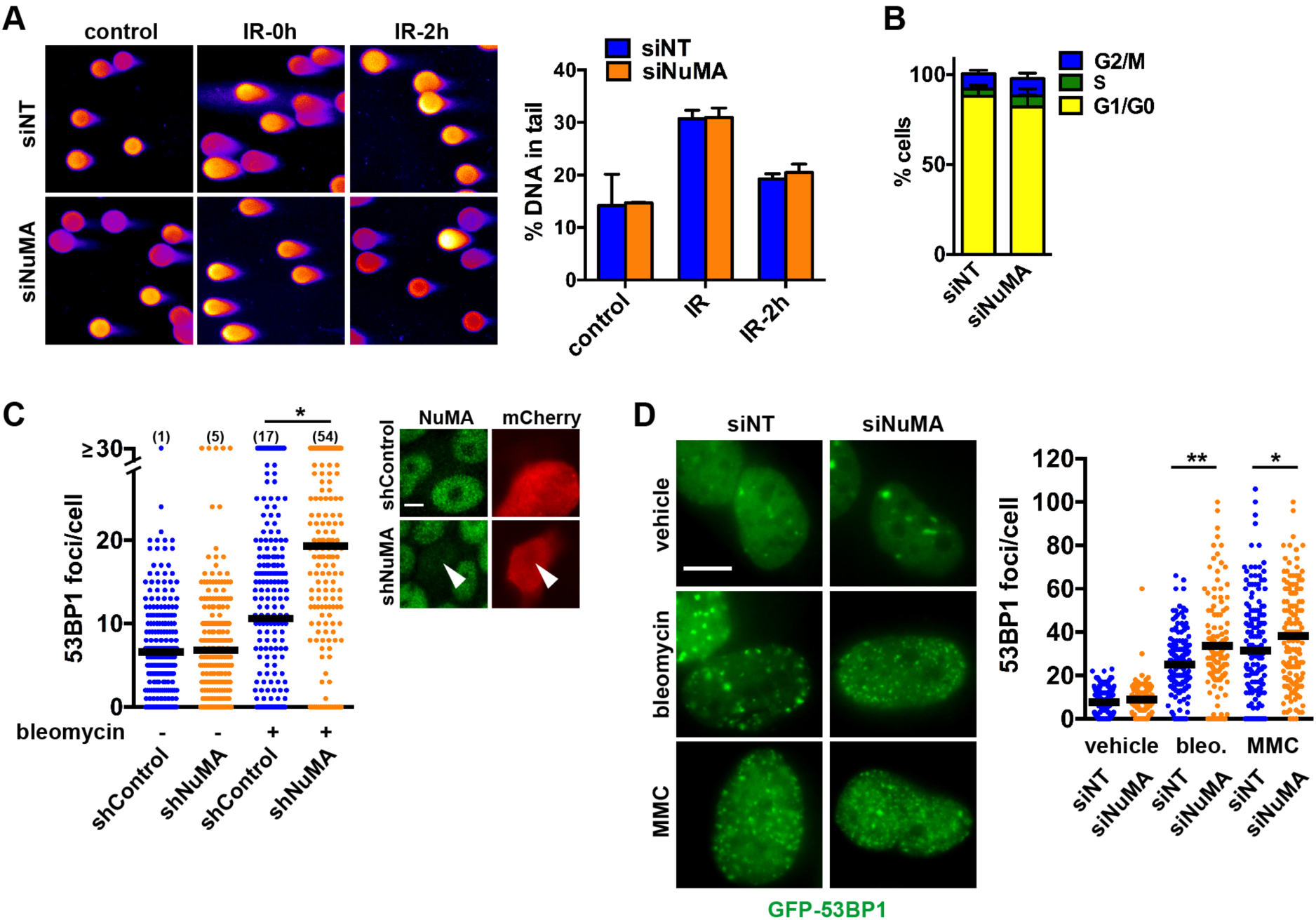
Accumulation of 53BP1 at DNA repair foci is impaired in cells silencing NuMA. **A** Detection of DSB with the neutral comet assay in HMT-3522 S1. Cells were transfected with nontargeting or NuMA siRNA, irradiated (3 Gy) and processed immediately or left to recover for 2h. Controls were mock irradiated. Average proportions of DNA in comet tails are shown in the bar graph (n = 3) **B** Cell cycle distribution in HMT-3522 S1 cells transfected with NuMA-specific siRNA or with nonspecific siRNA. **C** Enumeration of 53BP1 foci in HMT-3522 S1 cells transfected with constructs encoding NuMA specific (shNuMA) or scrambled (shControl) small hairpin RNA. Six days after transfection, cells were treated with BLM or vehicle. The number of cells with ≥30 53BP1 foci is indicated for each condition in parentheses. *, *P* < 0.001 (Kruskal-Wallis; n ≥ 180 cells from two biological replicates). mCherry was cotransfected with the shRNA to identify transfectants, as illustrated in the micrographs. **D** Representative GFP-53BP1 fluorescence images and GFP-53BP1 foci quantification (scatter plot) in U2OS cells transfected with nontargeting (NT) or NuMA siRNA. Cells were treated with vehicle, bleomycin, or MMC. Scale bar, 10 μm. *, *P* < 0.05 and **, *P* < 0.005 (ANOVA and Tukey; n > 100 cells from three biological replicates).

**Figure S4.**
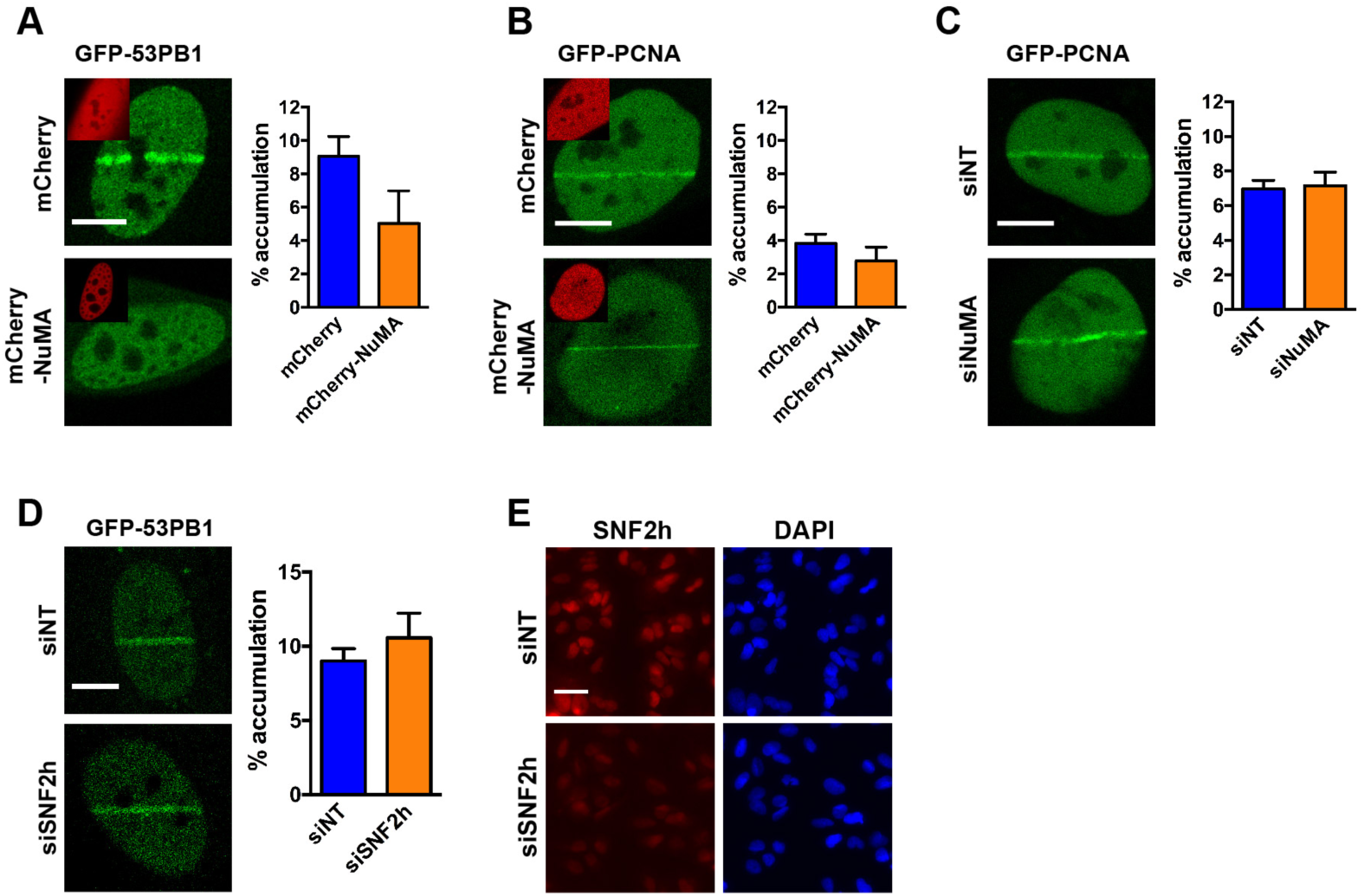
Inhibition of 53BP1 recruitment to DNA damage sites by NuMA is not mediated by SNF2h. **A** Accumulation of GPF-tagged 53BP1 at laser-microirradiated tracks in U2OS cells expressing mCherry or mCherry-NuMA. The graph represents the fractions of GFP signals at the tracks. **B-C** Accumulation of GFP-PCNA at laser-microirradiated tracks in cells overexpressing **(B)** or silencing **(C)** NuMA. **D** GFP-53BP1 accumulation at laser-microirradiated tracks in cells transfected with nontargeting siRNA or with SNF2h siRNA. Scale bars, 10 μm (A-D). **E** Verification of SNF2h silencing by fluorescence microscopy. Scale bar, 100 μm

**Figure S5.**
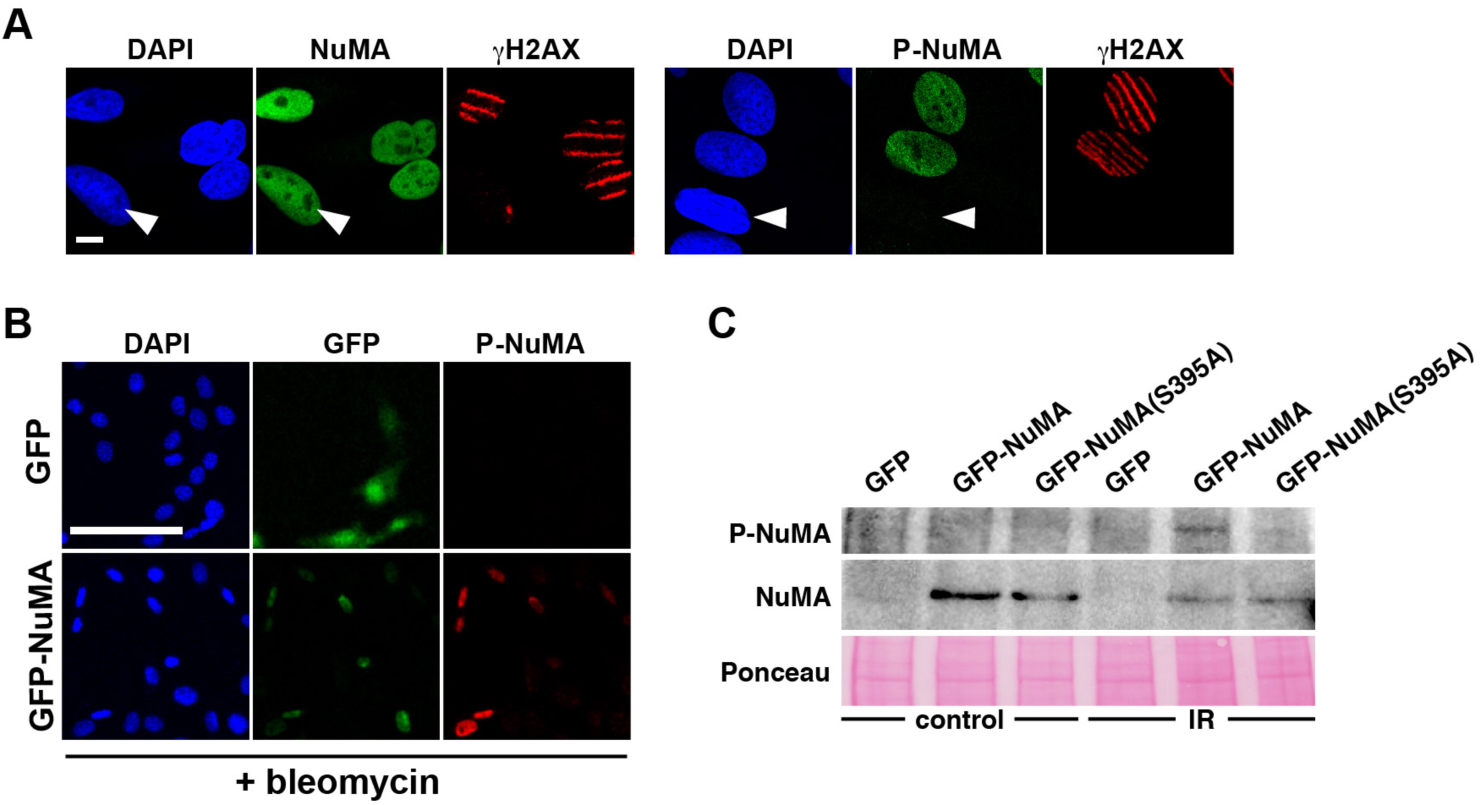
Pan-nuclear localization of P-NuMA. **A** Immunostaining of U2OS cells for NuMA (left) or P-NuMA (serine 395; right) after laser microirradiation. Arrowheads indicate nonirradiated cells, as evidenced by the lack of γH2AX signals. Scale, 10 μm. **B** Validation of P-NuMA detection by immunostaining in mouse NIH3T3 fibroblasts expressing GFP (top) or GFP-NuMA (human sequence; bottom) after bleomycin treatment (20 mU/ml; 1h). Scale, 100 μm. **C** Detection of NuMA and P-NuMA by immunoblot in NIH3T3 cell extracts. Cells were transfected with GFP, GFP-NuMA, or the nonphosphorytable mutant GFP-NuMA(S395A). Lanes on the right correspond to irradiated samples (IR, 10 Gy).

**Figure S6.**
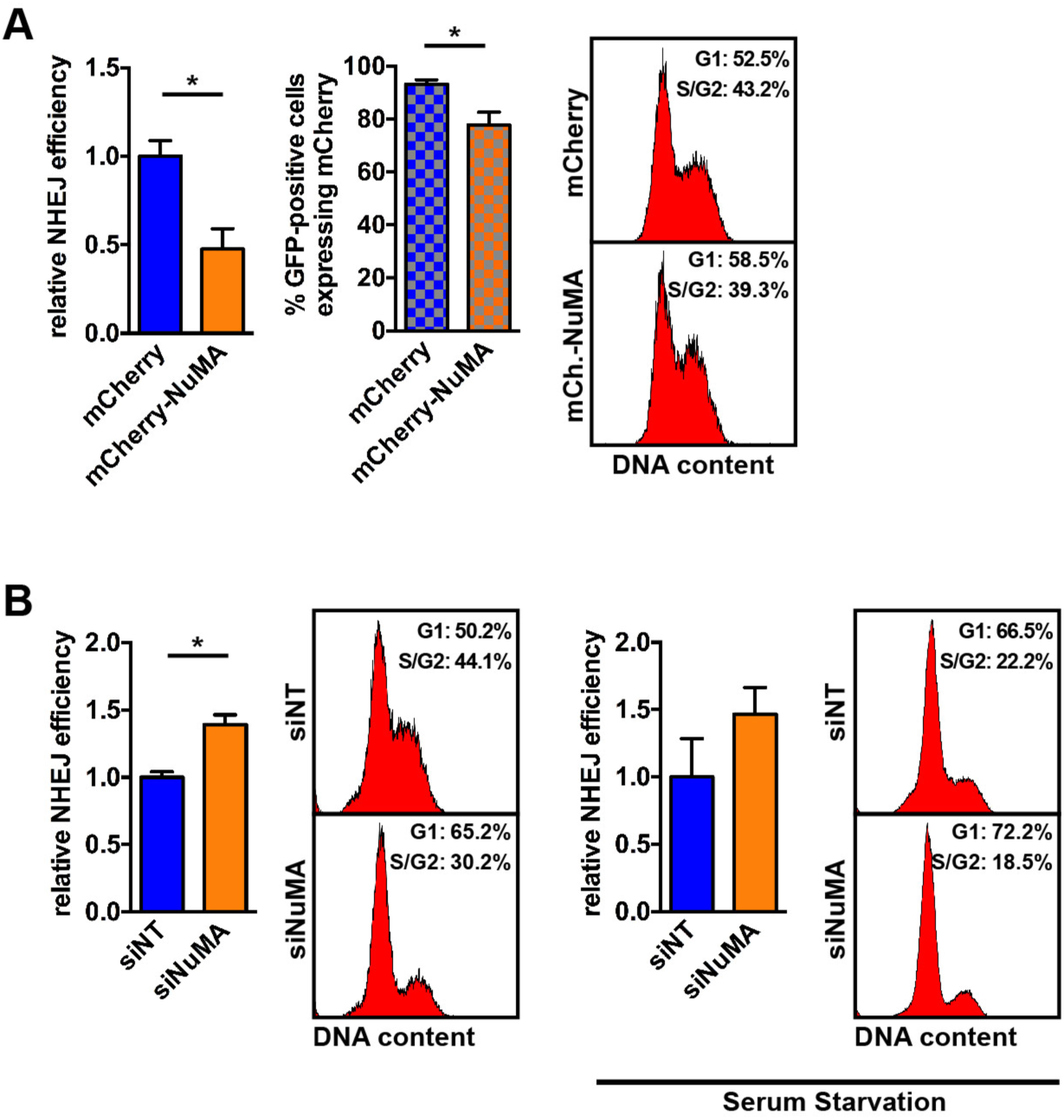
NuMA negatively regulates NHEJ. **A** Quantification of NHEJ activity in U2OS cells with a stably integrated NHEJ reporter constituted of the GFP coding sequence interrupted by an exogenous exon flanked by ISceI recognition sites. NHEJ reconstitutes the GFP coding sequence after ISceI cleavage. Normalized fractions of GFP-positive cells 24h after transfection with ISceI and mCherry or mCherry-NuMA are shown in the bar graph (left). The fraction of cells with mCherry fluorescence among GFP-expressing cells is shown in the cross-ruled graph whereas cell cycle distribution is shown on the right. **B** Quantification of repair events in U2OS-NHEJ cells transfected with NuMA or with nontargeting (NT) siRNA, and with ISceI, 48h and 24h before flow cytometry analysis, respectively. The same experiment was done in serum-starved cells to normalize cell cycle distribution. *, *P* < 0.05 (t-test, n = 4)

**Figure S7.**
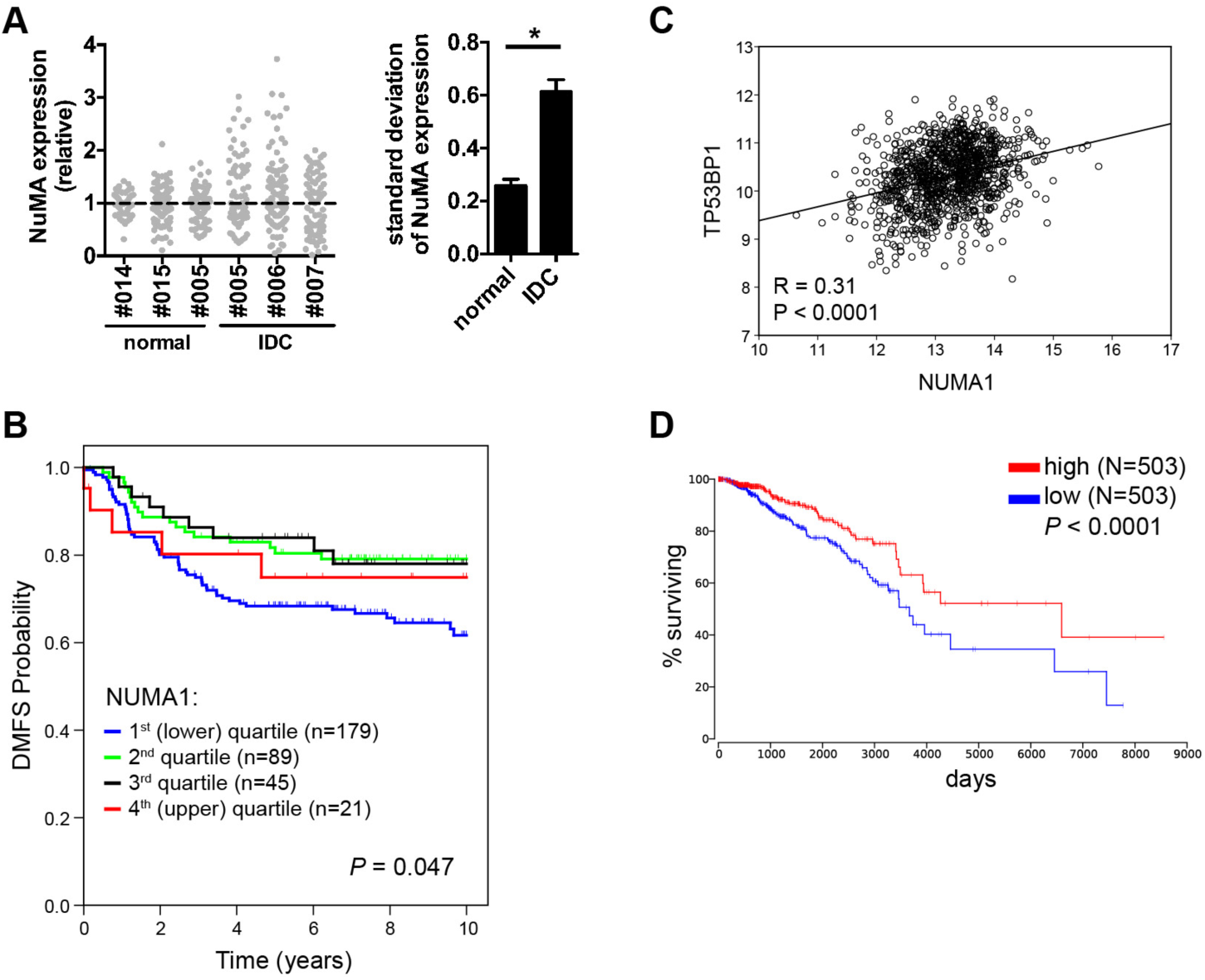
NuMA expression predicts survival in breast cancer patients. **A** NuMA expression in IDC (patients #005-7) and in normal tissues from reduction mammoplasties (#014; #015) or adjacent to a tumor (#005). NuMA staining intensities are normalized to the median value of the image. Each point from the scatter plot represents a cell nucleus. Standard deviations are shown in the bar graph. *, *P* < 0.005 (t-test). **B** Kaplan-Meier plot of distant metastasis-free survival (DMFS) based on *NUMA1* expression in a subset of patients with basal-like tumors (from (Nagalla, Chou et al., 2013)). The log-rank p-value is shown. **C** Correlation analysis of *NUMA1* and *TP53BP1* mRNA expression levels in 1,079 breast tumors of The Cancer Genome Atlas cohort (TCGA; data expressed as log2 RNA-seq normalized read counts). **D** Overall survival of patients from the TCGA cohort stratified by *NUMA1* expression above (red) vs. below (blue) median.

